# Modulation of cellular ageing regulates Integrated Stress Response signalling to control blood cell homeostasis

**DOI:** 10.1101/2025.01.09.632127

**Authors:** Kishalay Ghosh, Rohit Krishnan Iyer, Saloni Sood, Mohamed Sabeelil Islam, Jyotsana G. Labad, Rohan Jayant Khadilkar

## Abstract

Organismal ageing is associated with a decline in cellular and molecular functions. One of the hallmarks of ageing is stem cell exhaustion which occurs due to dynamic changes in the stem cell – niche microenvironment. Ageing affects stem cell potency impacting self-renewal and differentiation trajectories. Blood cell homeostasis in the hematopoietic system is tightly regulated by a fine balance between stemness and differentiation. Here, we employ the *Drosophila* hematopoietic organ, the larval lymph gland (LG) to investigate the impact of modulating cellular ageing on organ homeostasis. LG consists of a Posterior Signalling Center (PSC) that acts as a stem cell niche and maintains the prohemocytes housed in the Medullary Zone (MZ). Cortical Zone (CZ) consists of differentiated hemocytes namely plasmatocytes, crystal cells and lamellocytes. Intricate signalling networks active in the PSC and MZ orchestrate and regulate homeostasis. In this study, we characterize the effect of genetic perturbation of the molecular circuitry of ageing locally and systemically to investigate its effect on LG blood cell homeostasis. Genetic modulation of cellular ageing displays traditional hallmarks of ageing validating our model in the LG. Our results indicate that induction of accelerated ageing both locally and systemically leads to a reduction in stem cell niche size, DNA damage accumulation and increased progenitor differentiation whereas decelerated ageing shows an opposite trend. We show that the Integrated Stress Response (ISR) pathway is activated upon inducing accelerated ageing in the LG hemocytes possibly to recoup back to homeostatic conditions. Furthermore, LG’s of ISR pathway mutants or upon perturbation of ISR pathway components in prohemocytes show increased blood cell differentiation indicating disruption of homeostasis. Genetic epistasis analysis shows that ectopic over-expression of ISR pathway genes in an accelerated ageing scenario over and above the existing levels can rescue the defects in blood cell homeostasis. Overall, our study explores how modulation of cellular aging locally or systemically can impact tissue homeostasis. Our research paves way to understand the cellular mechanisms underlying an aged versus young stem cell-niche microenvironment and how its abrogation may lead to onset of disease.

## Introduction

Ageing is characterized by progressive decline in cellular and molecular functions that lead to loss of physiological integrity impacting organismal function. There are a number of physiological disorders like cancer, diabetes, cardiovascular and neurodegenerative disorders that are an outcome of risk factors closely associated with ageing (Carlos Lopez-Otin., 2023). The hallmarks of ageing represent qualitative and quantitative decline in cellular and molecular functions as age progresses like genomic instability, loss of proteostasis, mitochondrial dysfunction, stem cell exhaustion etc. which adversely affect organismal lifespan. Stemness refers to self-renewal ability of stem cells and their capability of multi-directional differentiation into other cell types that is gradually affected upon ageing (Yi et al., 2020). The functionality of stem cells is influenced by the microenvironment or niche in which they reside which serves as a spatial organization providing protection and enabling functional interactions among various cell types like immune cells, endothelial cells, mesenchymal cells etc. This stem cell microenvironment is rich in extracellular matrix (ECM), signaling molecules including growth factors and cytokines that influence stem cell behaviour impacting self-renewal and their ability to differentiate. Upon ageing, there are significant changes in the stem cell niche including cell-extrinsic factors like accumulation of inflammatory cytokines and decrease in growth factors promoting regeneration, alterations in the stem cell niche structure due to changes in ECM composition and cell-intrinsic factors like perturbation in protein homeostasis, mitochondrial dysfunction, accumulation of ROS and DNA damage, cellular senescence, epigenetic reprogramming, and so on (Farahzadi et al., 2023). One of the adult stem cell types that is known to be impacted by aging are the Hematopoietic Stem Cells (HSCs) that produce all mature blood cell types. As HSCs age, they show increased myeloid-biased differentiation at the expense of lymphopoiesis where myeloid differentiation linked genes were shown to be upregulated (Rossi et al., 2005). Another characteristic feature of ageing is remodelling of the bone marrow (BM) niche where HSCs relocate within the BM niche, away from the bone surface (endosteum) to the central regions of BM niche (Ho & Méndez-Ferrer, 2020). During ageing of HSCs, there is increased accumulation of pro-inflammatory cytokines including CC-chemokine ligand 5 (CCL5), impaired autophagy, adipocyte skewing, accumulation of DNA damage and ROS, inactivation of Drp1 (Dynamin related protein 1) mediating mitochondrial fission and enhanced mitochondrial oxidative phosphorylation (OXPHOS), all leading to decline in HSC stemness (Liu et al., 2022).

Age-related alterations occur in various mature blood cell types like B-cells, T-cells and hematopoietic stem and progenitor cells (HSPCs). Due to age-related gradual acquisition of somatic mutations or changes in the niche, there is reduced genetic diversity not only at the level of stem and progenitor cells but also mature blood cell populations. This occurs due to gradual clonal expansion of stem cells carrying mutations or genetic variants that are disruptive resulting in age-related clonal hematopoiesis (ARCH). ARCH treatment and prevention are a major challenge currently as it has a high risk of leading to hematological malignancies (Shlush, 2018). Due to these challenges associated with ARCH, a genetically tractable model to understand the mechanisms governing clonal hematopoiesis becomes important. Here, we use *Drosophila* as a model organism and its larval hematopoietic organ, the lymph gland (LG) to understand the effects of genetic modulation of cellular ageing in different blood cell subsets in *Drosophila* both locally and systemically.

*Drosophila* hematopoietic system serves as an excellent model due to its striking similarities with the mammalian system, availability of sophisticated genetic tools and experimental ease to understand molecular mechanisms governing hematopoiesis (Evans et al., 2003). Hematopoiesis in *Drosophila* occurs in two distinct waves: The first wave takes place during embryogenesis, where hematopoietic progenitors originate from the pro-cephalic head mesoderm (Evans et al., 2003). These progenitors differentiate into various hemocyte types, primarily plasmatocytes and crystal cells. Plasmatocytes comprising over 90% of hemocytes are functionally similar to mammalian macrophages, engaging in phagocytosis and immune responses. Crystal cells account for about 5% of hemocytes and are involved in wound healing, the synthesis of melanin through prophenoloxidase (PPO) and recently reported to be oxygen carriers (Shin et al. 2024). The second wave occurs during larval development, primarily within a specialized organ known as the lymph gland (Lanot et al., 2001, Jung et al., 2005). This organ serves as an active hub for hematopoietic activity, where progenitor cells maintain their population and upon receiving required cues differentiate into mature hemocytes. There is a third blood cell type in *Drosophila* named lamellocytes which differentiate from prohemocytes or trans-differentiate from other hemocyte types under acute stress like wasp infestation (Rizki et al., 1979, Franc et al., 1996). The lymph gland is organized into distinct zones: a Posterior Signaling Center (PSC) which acts as a niche regulating both self-renewal and differentiation of prohemocytes, medullary zone (MZ) that houses quiescent prohemocytes, an intermediate zone (IZ) that contains hemocytes transitioning towards mature differentiated cells and cortical zone (CZ) for mature differentiated hemocytes namely plasmatocytes, crystal cells and lamellocytes (Jung et al., 2005, Kharrat et al., 2022). Recent single cell RNA sequencing studies demonstrated a considerable transcriptomic and functional heterogeneity within these broadly classified compartments for example we now know that the progenitors in MZ could be either core or distal hematopoietic progenitors (Fu et al., 2020, Cattenoz et al., 2020, Tattikota et al., 2020, Cho et al., 2020, Girard et al., 2021, Kharrat et al., 2022). Multiple signalling pathways play an important role not only in maintaining the PSC niche population but also in regulating the differentiation of the prohemocytes that reside in the MZ. Prohemocytes in the MZ are maintained by Hedgehog and Serrate mediated Notch signalling from the PSC (Lebetsky et al., 2003, Mandal et al., 2007, Baldeosingh et al., 2018). Signalling pathways like wingless (Sinenko et al., 2009, Goins et al., 2024), Dpp (Pennetier et al., 2012, Dey et al., 2016), JAK/STAT (Minakhina et al., 2011, Sinha et al., 2013, Rodrigues et al., 2021) also regulate prohemocytes in the MZ. Such tight regulation maintains a balance between stem cell quiescence and activation, ensuring a steady supply of progenitors for differentiation into various hemocyte types (Duvic et al., 2002, Benmimoun et al. 2012, Pennetier et al., 2012, Banerjee et al., 2019, Kharrat et al., 2022). Similarly, a complex interplay of different LG intrinsic and extrinsic signaling pathways determine the extent of differentiation and dictate the lineage commitment decision for different blood cell types (Koranteng et al., 2022). For example, Pvr (PDGF/VEGF), EGFR promotes plasmatocyte differentiation while Notch and Hippo signaling are crucial for crystal cell differentiation (Milton et al., 2014, Dey et al, 2016, Khadilkar and Tanentzapf, 2019, Zhang et al. 2023). Under stressful conditions like increased ROS levels, injury, infection or wasp infestation, signals like JNK, Toll, EGFR pathway are activated to promote lamellocyte differentiation, a hemocyte type found only under such situations (Meister and Ferrandon, 2011; Anderl et al., 2016, Louradour et al., 2017). The complexity and heterogeneity of cell subsets in the LG coupled with the genetic tools available for gene manipulation in a spatial and temporal manner in the hematopoietic system in *Drosophila* provide a scorable system to assess the impact of genetically modulating cellular ageing on blood cell homeostasis.

Now, to understand if genetic modulation of cellular ageing has any impact on stem cell and tissue homeostasis we used two different approaches – one for accelerating ageing and the other for decelerating ageing utilizing the *Drosophila* larval lymph gland as our model. We use the “inflammaging” approach to induce accelerated ageing which refers to chronic and persistent activation of inflammation. It negatively influences overall lifespan and alters stem cell function by promoting a chronic hyperactive immune state with increased expression of anti-microbial peptides & pro-inflammatory cytokines leading to cellular senescence and impaired stem cell regeneration in *Drosophila* (Garschall and Flatt., 2018, Zhang et al., 2021). We activate the classical NF-κB signalling pathways – Toll and Imd in *Drosophila* to induce accelerated ageing. Pirk (peptidoglycan recognition protein (PGRP)-interacting Receptor Kinase) acts as a negative feedback regulator of Imd (Immune deficiency) pathway which is a component of NF-κB signaling cascade and involved in eliciting immune response against Gram-negative bacteria in the form of anti-microbial peptides (AMPs) like Diptericin, Drosocin etc. (Kounatidis et al., 2021). Pirk interacts with Imd directly and the cytoplasmic tail of PGRP-lysine type (LC) and forms a complex that inactivates them (Pechmann et al., 2017) and prevents nuclear localisation of Relish, a NF-κB transcription factor, thereby regulating and aiding in eliciting a controlled Imd immune response against pathogens (Kaur et al., 2020). Knockdown of *pirk* leads to upregulation of Imd pathway (Badinloo et al., 2021) and in staging a chronic and persistent inflammatory state that systemically accelerates ageing and reduces lifespan at an organismal level (Giannakou & Partridge, 2007) by inducing neurodegenerative conditions and metabolic alterations (Yamashita et al., 2021) including altered lipid metabolism due to depleting glycogen reserves in fat body and increased circulation of sugar as response to infection and inflammation (Davoodi et al., 2019). Apart from Imd activation, we target the Toll pathway. When there is heightened activation of the Toll pathway leading to elevated AMP expression and decreased immune efficiency (immunosenescence), it corresponds to reduced lifespan systemically at an organismal level (Kounatidis et al., 2020). Toll10B is a gain-of-function mutation of Toll receptor that constitutively activates the Toll pathway (Artero et al., 2003) and thus we have used it for induction of inflammaging via Toll. For inducing decelerated cellular ageing, we employ the over-expression of Foxo or Atg8. Foxo (Forkhead box O) are proteins under the Forkhead family of transcription factors and characterized by a conserved DNA-binding domain known as Forkhead box (FOX). They aid in lifespan extension by decreasing insulin-like growth factor (IGF)-like signaling (IIS) and inducing a state similar to caloric restriction (Broughton et al., 2005), enhancing autophagy resulting in the clearance of damaged and accumulated proteins and organelles (Morselli et al., 2010), upregulating stress-responsive genes and down-regulating genes of oxidative phosphorylation to tackle oxidative and metabolic stress (Hwangbo et al., 2004). Studies have shown that over-expressing dFoxO in *Drosophila* fat body and gut under specific dietary conditions extended longevity as compared to the *wildtype* control (Broughton et al., 2005). In *C. elegans*, activation of DAF-16, a FoxO homolog, promoted oxidative stress resistance and enhanced autophagy, thereby extending lifespan (Kenyon, 2010). *Drosophila* Atg8a is an autophagic protein with a crucial role in autophagosome biogenesis and maturation which is important for degradation and clearance of damaged and accumulated cellular components, thereby maintaining cellular homeostasis, improving metabolic efficiency and extending lifespan (Yin et al., 2016). Also, it binds to transcription factors like Sequoia that regulates autophagy gene expression (Jacomin et al., 2019). Atg8a also interacts with Sirtuins which are upregulated in conditions of nutrient stress which then deacetylate Atg8a to activate autophagy under starvation conditions (Jacomin et al., 2020). Additionally, it was demonstrated that over-expressing Atg8a in specific tissues like muscles and gut resulted in enhanced tissue integrity and function and systemically led to an overall organismal longevity and improved health span (Bai et al., 2013, Li et al., 2021). Atg8a is also known to promote mitochondrial function by upregulating genes involved in mitochondrial biogenesis and mitochondria-targeted mitophagy that degrades damaged and impaired mitochondria contributing to high ROS levels and improves the overall quality of mitochondrial population within cells (Youle & van der Bliek, 2012).

In this study, we have done a detailed characterization of how tissue homeostasis in the *Drosophila* LG is impacted upon genetic modulation of cellular ageing not only by genetic perturbations locally in different cell subsets of the LG but also systemically. We validate these genetic modalities and also use alternate approaches including chemical or drug-based interventions for accelerating or decelerating ageing and score for homeostasis. Furthermore, we uncover signalling mechanism that kicks in when the cells in the LG sense stress induced by accelerated ageing and our results show that the ectopic Integrated Stress Response (ISR) pathway activation over and above its existing levels are capable of restoring blood cell homeostasis in the LG when it is undergoing accelerated cellular ageing. Our results provide important insights into the biology of ageing of the stem cell niche micro-environment which often transforms into a micro-environment similar to a leukemogenic niche that fosters malignancies in such adverse conditions.

## Results

### Lymph gland hematopoietic progenitors display typical hallmarks of aging upon genetic modulation of cellular aging

Ageing is associated with a progressive decline in cellular and molecular functions along with increase in mortality (Silva & Schumacher., 2021). Here, we set out to understand the effect of modulating the molecular circuitry of aging in various cellular subsets of the *Drosophila* larval lymph gland to study the effects on stem cell and tissue homeostasis. We started by functionally validating our cellular ageing models – both accelerated and decelerated ageing. Here, we have modulated the molecular circuitry of aging in the blood progenitors in the lymph gland. Induction of accelerated ageing was achieved by upregulation of NF-***κ***B signaling pathways like Imd activation via *pirk* knockdown or Toll pathway over-activation. *pirk* is a negative regulator of *Drosophila* Imd pathway and RNAi-mediated knockdown of *pirk* causes Imd hyperactivation (Kleino et al., 2008). Toll pathway was activated via over-expression of Toll10B, a gain of function and constitutively active Toll receptor mutant whose over-expression leads to hyperactivation of Toll pathway (Schmid et al., 2014). Decelerated aging was induced through ectopic over-expression of Atg8, an important autophagy-related protein required for autophagosome biogenesis and maturation and induction of autophagy (Nair et al., 2012) & Foxo, a transcription factor regulating diverse biological processes including aging and metabolism (Du & Zheng., 2021). We employed the above transgenes for modulation of cellular aging as previous literature suggested that Imd or Toll pathway upregulation predisposed fly brains to toxic levels of AMPs, increased inflammation and dramatically reduced lifespan (Kounatidis et al., 2017; Khor & Cai., 2020) and over-expression of Atg8 in neurons or Foxo over-expression in the pericerebral fat body led to an increase in lifespan at an organismal level in *Drosophila* (Gelino & Hansen., 2013; Du & Zheng., 2021). In our study, we have assessed the typical hallmarks of cellular aging including regulation of autophagic flux, reactive oxygen species (ROS) levels and regulation of proteostasis i.e. protein turnover assessment (Carlos López-Otín et al., 2023) upon genetic modulation of cellular aging in the *tepIV*-positive core progenitors of lymph gland.

For estimation of autophagic flux we scored for p62, an adaptor protein that mediates interaction between the cargo and Atg8 on autophagosomes for cargo degradation and gets cleared along with the cargo during autophagy (Bjørkøy et al., 2009). We have estimated the ratio of p62 positive puncta per cell upon modulation of ageing and our results indicate that there are decreased p62 positive puncta upon over-expression of Atg8 in *tepIV -* positive progenitors (Fig. 1B, D) whereas an accumulation of p62 upon Imd activation (Fig. 1C, D) as compared to the *wildtype* control (Fig. 1A, D). These results indicate that autophagy in the blood progenitors is impaired upon induction of accelerated ageing whereas in decelerated aging scenario, there is degradation of p62 moiety along with the autophagic cargo suggesting an efficient and functional autophagy process. ROS levels measured as mean fluorescence intensity levels were not altered upon induction of decelerated aging via Atg8 or Foxo over-expression (Fig. 1F, G, I) while Toll over-activation in progenitors exhibited accumulation of ROS (Fig. 1H – I) as compared to *wildtype* control (Fig. 1E, I). Similarly for the assessment of regulation of proteostasis, protein turnover was estimated using Proteostat reagent (Basisty et al., 2018, Kakraba et al., 2023) and we find that *pirk* knockdown or Toll pathway upregulation showed significant protein aggregate accumulation (Fig. 1M-O) indicating a net decrease in protein turnover and loss of proteostasis whereas decelerated ageing induced by Atg8 or Foxo over-expression displayed significant suppression of protein aggregate formation (Fig. 1K, L, O) indicating increase in protein turnover and restoration of protein homeostasis as compared to *wildtype* control (1J, O).

**Figure 1:**
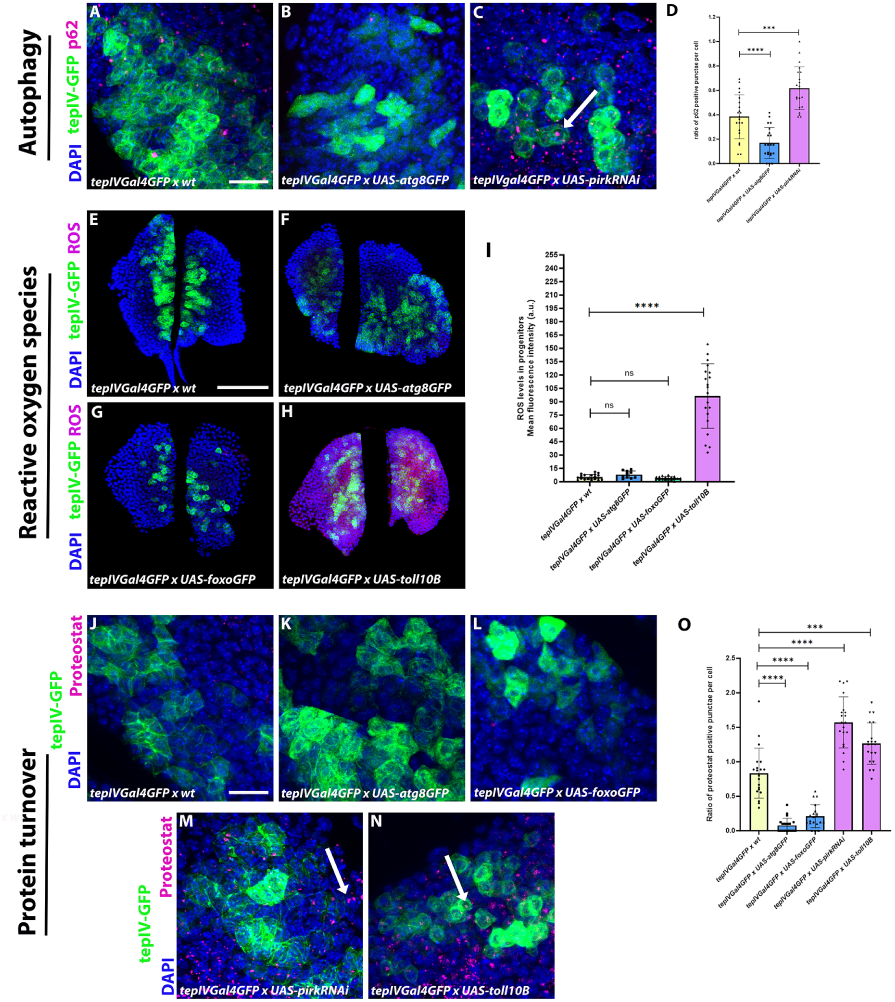
Validation of the hallmarks of aging upon modulation of cellular aging in hematopoietic progenitors of the lymph gland. Estimation of autophagic flux by evaluating the ratio of p62 positive punctae per cell marked by p62 antibody (magenta) upon core progenitor-specific (using *tepIVGal4*) knockdown of *pirk* or over-expression of *atg8* as compared to *wildtype* control (Fig. 1A-D). Mean fluorescence intensity of ROS levels in the core progenitors marked by Cell-ROX deep red reagent (Red) upon *tepIVGal4* mediated expression of *UAS-atg8GFP, UAS-foxoGFP* or *UAS-toll10B* as compared to *wildtype* control (Fig. 1E-I). Estimation of protein turnover by estimating the ratio of Proteostat positive punctae per cell marked by PROTEOSTAT detection reagent (magenta) upon *tepIVGal4* mediated expression of *UAS-atg8GFP, UAS-foxoGFP, UAS-pirkRNAi* or *UAS-toll10B* as compared to *wildtype* control (Fig. 1J-O). Graphical representation of the ratio of p62 positive punctae per cell (Fig. 1D) or Mean fluorescence intensity of ROS levels in the *tepIV*-positive core progenitors (Fig. 1I) or ratio of Proteostat positive punctae per cell (Fig. 1O) upon expression of *UAS-atg8GFP, UAS-foxoGFP, UAS-pirkRNAi* or *UAS-toll10B* in the progenitors as compared to *wildtype* control. For autophagic flux estimation by p62 staining, *tepIVGal4 x UAS-pirkRNAi* (N = 11, n = 21) and *tepIVGal4 x UAS-atg8GFP* (N = 11, n = 21) were analyzed as compared to *wildtype* control, *tepIVGal4 x wt* (N = 11, n = 21). For estimation of ROS levels by Cell-ROX deep red staining, *tepIVGal4 x UAS-atg8GFP* (N = 5, n = 10), *tepIVGal4 x UAS-foxoGFP* (N = 9, n = 18) and *tepIVGal4 x UAS-toll10B* (N = 10, n = 20) were analyzed as compared to *wildtype* control, *tepIVGal4 x wt* (N = 10, n = 20). For estimation of protein turnover by Proteostat detection reagent staining, *tepIVGal4 x UAS-atg8GFP* (N = 10, n = 20), *tepIVGal4 x UAS-foxoGFP* (N = 10, n = 20), *tepIVGal4 x UAS-pirkRNAi* (N = 10, n = 20) and *tepIVGal4 x UAS-toll10B* (N = 10, n = 20) were analyzed as compared to *tepIVGal4 x wt* (N = 10, n = 20) *as wildtype control*. N denotes the number of larvae & n denotes the number of individual primary lymph gland lobes analyzed per genotype. Individual data points in the graphs represent individual primary lobes of the Lymph gland. GFP expression (green) is driven by *tepIVGal4* (Fig. 1A-1C, 1E-1H, 1J-1N). Nuclei are stained with DAPI (Blue). Values are mean ± SD, and asterisks denote statistically significant differences (ns denotes not significant, *** p < 0.001, ****p < 0.0001). Student’s t-test with Welch’s correction was performed for the statistical analysis. Scale Bar: 30 µm (A-C, J-N), 50 µm (E-H).

### Hematopoietic niche size in the lymph gland is altered upon localized or systemic perturbation of cellular ageing

Posterior Signaling Centre (PSC) serves as the hematopoietic progenitor-niche or microenvironment that interacts with the hematopoietic progenitors through signaling to maintain a balance between progenitor maintenance and differentiation (Krzemień et al., 2007, Mandal et al., 2007, Pennetier et al., 2012). Here, we investigate the effect of aging modulation on the size of PSC niche in lymph gland that regulates blood cell homeostasis in *Drosophila*. Previous studies have shown that downregulation of the Imd pathway by Relish depletion exhibited PSC niche hyperplasia and aberrant differentiation of progenitors (Ramesh et al., 2021). We first performed PSC- niche specific induction of accelerated or decelerated ageing and our results indicated that over-activation of Toll using niche- specific *collierGal4* leads to a decrease in Antennapedia positive niche cells (Fig. 2E, a) whereas Imd activation has no effect on the niche (Fig. 2D, a). Niche- specific over-expression of Atg8 has no significant effect on PSC size (Fig. 2B, b) whereas Foxo over-expression leads to a decrease in niche size (Fig. 2C, b) as compared to the *wildtype* (Fig. 2A, a and b). We then assessed if there is any cell non-autonomous effect of progenitor- specific ageing modulation using *domeGal4* on the niche size. Similar to the cell-autonomous effect of Toll activation, progenitor-specific Toll activation resulted in a decrease in niche size whereas Imd activation did not alter niche size as compared to the *wildtype* (Fig. 2F, I, J and c). Decelerated ageing induction in the progenitors had no effect on niche size as compared to the *wildtype* (Fig. 2F, G, H and d). Previous studies have shown that differentiated hemocytes in the LG can reciprocally regulate the blood progenitors (Mondal et al., 2011) however whether the cortical zone hemocytes regulate niche size is underexplored. We hypothesized that modulating ageing in mature differentiated hemocytes could remodel the niche in the organ. In order to test this, we used *hmlΔGal4* that drives expression in cortical zone differentiated hemocytes. *hmlΔGal4* mediated induction of accelerated or decelerated ageing has no effect on the niche size as compared to the *wildtype* (Fig. 2K-O, e and f). We then investigated the effect of modulating ageing in the whole lymph gland using *e33cGal4* and found that accelerating ageing by both Toll or Imd activation resulted in a smaller niche (Fig. 2S, T and g) whereas induction of decelerated ageing by Atg8 over-expression had no effect on the niche (Fig. 2Q and h). Foxo over-expression very similar to its cell autonomous effect on the niche led to a decrease in niche size upon LG-specific over-expression (Fig. 2R and h) as compared to *wildtype* control (Fig. 2P, g and h). Since lymph gland cells can respond to systemic changes as documented earlier (Shim et al., 2012, Cho et al., 2018, Goyal et al., 2021, Koranteng et al., 2022), we checked if fat body specific modulation of ageing that is known to also impact overall lifespan of flies can alter the niche in the LG. We activated accelerated ageing in the fat body using *pplGal4* and it showed no alteration in LG niche size (Fig. 2X, Y and i) whereas Atg8 over-expression again had no effect on niche size (Fig. 2V and j) and Foxo led to a decrease in niche size (Fig. 2W and j) as compared to *wildtype* (Fig. 2U, i and j) indicating that perturbation of systemic insulin signalling exerts a long range effect on the LG niche size. Interestingly in the case of Foxo, a downstream target of Insulin pathway (Piug et al., 2005) that exerts feedback regulation of the pathway attenuating it (Ni et al., 2007) we find that cell-autonomous, whole LG-specific as well as systemic Foxo over-expression leads to a decrease in niche size demonstrating a negative impact on the insulin signalling that maintains the niche size (Benmimoun et al., 2012). Overall, our results indicate that induction of accelerated ageing decreases the niche size and decelerated ageing mediated by autophagy activation does not alter the niche size in both localized or systemic manner.

**Figure 2:**
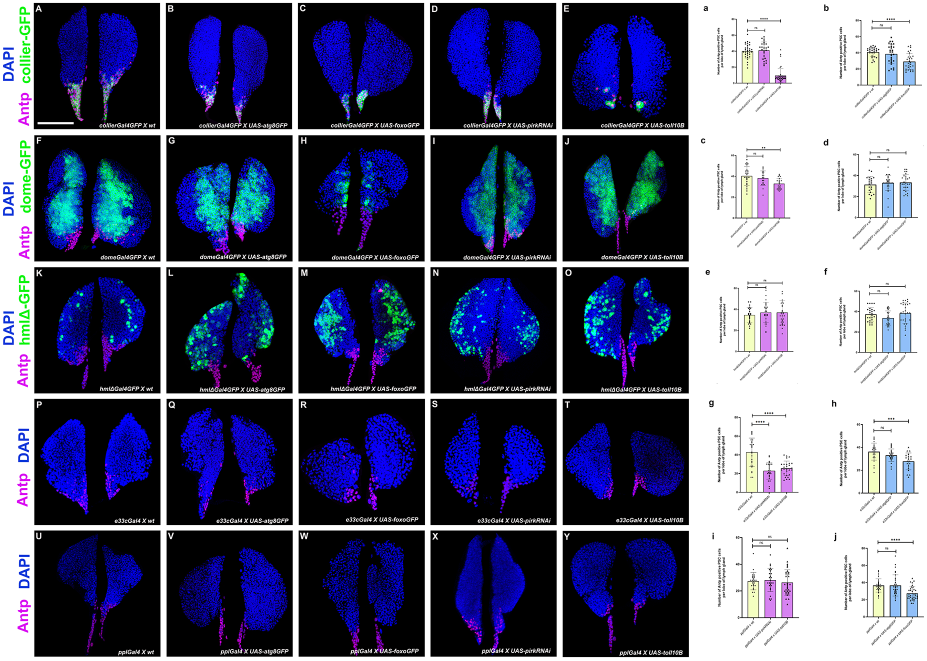
Localized perturbation in LG or systemic modulation of cellular ageing affects PSC niche size in *Drosophila*. Posterior Signalling Centre (PSC) niche cell population marked by Antennapedia (magenta) and graphical representation of niche cell numbers upon either PSC- specific (using *collierGal4*) or hematopoietic progenitor- specific (using *domeGal4*) or cortical zone differentiated hemocytes- specific (using *hmlΔGal4*) or whole lymph- specific (using *e33cGal4*) or systemic fat body- specific (using *pplGal4*) expression of *UAS-atg8GFP* (2B-V, 2b-j), *UAS-foxoGFP* (2C-W, 2b-j), *UAS-pirkRNAi* (2D-X, 2a-i) or *UAS-toll10B* (2E-Y, 2a-i) as compared to respective *wildtype* control (2A-U, 2a-j). For PSC niche - specific genetic modulation, *collierGal4 x UAS-atg8GFP* (N =14, n = 27), *collierGal4 x UAS-foxoGFP* (N = 15, n = 29), *collierGal4 x UAS-pirkRNAi* (N = 13, n = 25) and *collierGal4 x UAS-toll10B* (N = 20, n = 40) were analyzed as compared to *collierGal4 x wt* (N = 17, n = 33). For hematopoietic progenitor - specific genetic modulation, *domeGal4 x UAS-atg8GFP* (N = 14, n = 28), *domeGal4 x UAS-foxoGFP* (N = 14, n = 28), *domeGal4 x UAS-pirkRNAi* (N = 11, n = 21) and *domeGal4 x UAS-toll10B* (N = 11, n = 22) were analyzed as compared to *domeGal4 x wt* (N =12, n = 23). For differentiated hemocyte – specific genetic modulation, *hmlΔGal4 x UAS-atg8GFP* (N = 15, n = 30), *hmlΔGal4 x UAS-foxoGFP* (N = 15, n = 29), *hmlΔGal4 x UAS-pirkRNAi* (N = 14, n = 28) and *hmlΔGal4 x UAS-toll10B* (N = 16, n = 32) were analyzed as compared to *hmlΔGal4 x wt* (N = 14, n = 28). For whole lymph gland – specific genetic modulation, *e33cGal4 x UAS-atg8GFP* (N = 12, n = 23), *e33cGal4 x UAS-foxoGFP* (N = 10, n = 20), *e33cGal4 x UAS-pirkRNAi* (N = 12, n = 23) and *e33cGal4 x UAS-toll10B* (N = 14, n = 27) were analyzed as compared to *e33cGal4 x wt* (N = 14, n = 27). For fat body – specific genetic modulation, *pplGal4 x UAS-atg8GFP* (N = 15, n = 309), *pplGal4 x UAS-foxoGFP* (N = 17, n = 33), *pplGal4 x UAS-pirkRNAi* (N = 13, n = 26) and *pplGal4 x UAS-toll10B* (N = 16, n = 32) were analyzed as compared to *pplGal4 x wt* (N = 16, n = 32). Individual data points in the graphs represent individual primary lobes of the Lymph Gland. Nuclei are marked with DAPI (Blue). GFP is either driven by *collierGal4* (A-E), *domeGal4* (F-J) or *hmlΔGal4* (K-O). Values are mean ± SD, and asterisks denote statistically significant differences (ns denotes not significant, **p < 0.01, *** p < 0.001, ****p < 0.0001). Student’s t-test with Welch’s correction was performed for the statistical analysis. Scale Bar: 50 µm (A-Y).

### Modulation of cellular aging in the PSC niche regulates hemocyte differentiation in the lymph gland

Since modulating cellular ageing in a localized and systemic manner affected the PSC niche size, we wanted to determine if PSC- specific modulation of ageing impacted blood cell differentiation in the lymph gland. PSC-specific modulation of ageing was mediated by *collierGal4*. Niche specific activation of Toll or Imd pathway resulted in increased plasmatocyte differentiation (Fig. 3D, E and P) whereas over-expression of Atg8 did not affect plasmatocyte differentiation (Fig. 3B and Q). On the other hand, over-expression of Foxo in the niche which can attenuate insulin signaling increased plasmatocyte differentiation in the LG (Fig. 3C and Q) as compared to the *wildtype* control (Fig. 3A, P and Q). Induction of accelerated ageing in the niche by Imd activation triggered crystal cell differentiation (Fig. 3I, R) whereas Toll pathway activation resulted in a decrease in crystal cells (Fig. 1J, R). Induction of decelerated ageing especially by Foxo over-expression resulted in a decrease in crystal cell differentiation (Fig. 3H, S) and Atg8 over-expression had no significant effect (Fig. 3G, S) as compared to the control (Fig. 3F, R and S). Accelerated ageing induced by both Toll or Imd produced lymph glands positive for lamellocytes (Fig. 3N-O’, T) whereas Foxo over-expression in the niche led to a modest 15 % lymph glands being positive for lamellocytes (Fig. 3M-M’, T) and Atg8 over-expression caused no induction of lamellocyte differentiation (Fig. 3L-L’, T) as compared to the *wildtype* (Fig. 3K-K’, T). Previous studies have shown that wasp parasitism increases ROS levels activating Toll and EGFR pathway in the PSC which promotes aberrant progenitor differentiation into plasmatocytes and lamellocytes (Louradour et al., 2017) and trans-differentiation of circulating plasmatocytes into lamellocytes directly on the parasitic wasp eggs via increased Spitz secretion and EGFR signaling (Meister and Ferrandon, 2011; Anderl et al., 2016). The above literature supports our observation of elevated plasmatocyte and lamellocyte differentiation upon Toll pathway activation in the PSC niche.

**Figure 3:**
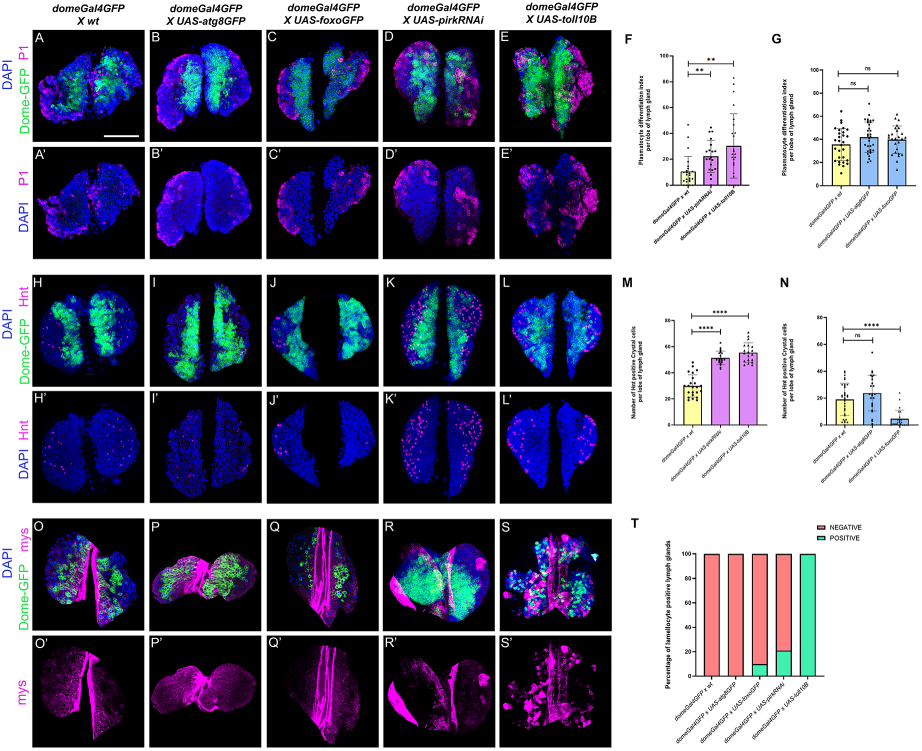
Localized distal progenitor subset-specific modulation of cellular ageing regulates LG homeostasis in *Drosophila*. Plasmatocyte differentiation marked by P1 (magenta) or crystal cell differentiation marked by Hindsight (Hnt) (magenta) or lamellocyte differentiation marked by β-integrin-specific Myospheroid (Mys) (magenta) upon progenitor-specific (using *domeGal4*) mediated expression of *UAS-atg8GFP* (Fig. 4B, 4I, 4P), *UAS-foxoGFP* (Fig. 3C, J, Q), *UAS-pirkRNAi* (Fig. 3D, K, R) or *UAS-toll10B* (Fig. 3E, L, S) as compared to *wildtype* control (Fig. 3A, H, O). Graphical representation of plasmatocyte differentiation index or number of crystal cells or percentage of lamellocyte positive lymph glands upon *domeGal4* mediated expression of *UAS-pirkRNAi* or *UAS-toll10B* (Fig. 3F, M, T) or *UAS-atg8GFP* or *UAS-foxoGFP* (Fig. 3G, N, T) as compared to *wildtype* control. For Plasmatocyte differentiation in accelerated ageing scenario: *domeGal4 x UAS-pirkRNAi* (N = 10, n = 20) and *domeGal4 x UAS-toll10B* (N = 12, n = 24) as compared to *domeGal4 x wt* (N = 12, n = 24) and in decelerated ageing scenario: *domeGal4 x UAS-atg8GFP* (N = 17, n = 33) and *domeGal4 x UAS-foxoGFP* (N = 13, n = 26) as compared to *domeGal4 x wt* (N = 14, n = 29). For crystal cell numbers in accelerated ageing scenario: *domeGal4 x UAS-pirkRNAi* (N = 11, n = 21) and *domeGal4 x UAS-toll10B* (N = 10, n = 19) as compared to *domeGal4 x wt* (N = 12, n = 23) and in decelerated ageing scenario: *domeGal4 x UAS-atg8GFP* (N = 12, n = 24) and *domeGal4 x UAS-foxoGFP* (N = 12, n = 23) as compared to *domeGal4 x wt* (N = 17, n = 33). N denotes the number of larvae & n denotes the number of individual primary lymph gland lobes analysed per genotype. Individual data points in the graphs represent individual primary lobes of the LG. GFP expression (green) is driven by *domeGal4* (Fig. 3A-S). Nuclei are stained with DAPI (Blue). Values are mean ± SD, and asterisks denote statistically significant differences (ns denotes not significant, **p<0.01, *** p<0.001, ****p<0.0001). Student’s t-test with Welch’s correction was performed for the statistical analysis. Scale Bar: 50 µm (A-S’).

### Hematopoietic progenitor- specific modulation of cellular ageing regulates blood cell homeostasis in *Drosophila*

Hematopoietic progenitors in the medullary zone (MZ) are important cellular subsets that form the basis of tissue homeostasis in the lymph gland. Hematopoietic progenitors not only respond to localized signals like Wg (Sinenko et al., 2009, Goins et al., 2024), Hh (Mandal et al., 2007, Dey et al., 2016, Baldeosingh et al., 2018), DPP (Pennetier et al., 2012, Dey et al., 2016), JAK-STAT (Minakhina et al., 2011, Sinha et al., 2013, Rodrigues et al., 2021) etc. for their maintenance but are also capable of sensing systemic signals (Benmimoun et al., 2012, Shim et al., 2012, Shim et al., 2013, Cho et al., 2018, Koranteng et al., 2022). Since our results indicated that the hematopoietic niche is impacted upon genetic modulation of cellular ageing, we set out to ascertain the impact of cellular ageing on the progenitors itself. Here, we focused on inducing accelerated or decelerated ageing in *Domeless*-positive distal progenitor population and *Chiz*-positive intermediate progenitors in the LG. Induction of accelerated ageing by Toll or Imd activation mediated by *domeGal4* led to an increase in both plasmatocyte and crystal cell differentiation (Fig. 3D-E’, K-L’ F, M) whereas induction of decelerated ageing resulted in no significant effect on plasmatocyte differentiation whereas crystal cell differentiation was suppressed in the case of Foxo over-expression (Fig. 3B-C’, I-J’, G, N) using *domeGal4* as compared to the *wildtype* (Fig. 3A-A’, H-H’, F, G, M and N). Also, Toll activation using *domeGal4* induced lamellocytes consistently in all lymph glands (100%) (Fig. 3S-S’, T) analyzed while Imd activation resulted in 20% lymph glands with lamellocytes (Fig. 3R-R’, T). In the case of decelerated ageing, Atg8 over-expression did not give any lymph glands with lamellocytes (Fig. 3P-P’, T) whereas Foxo over-expression resulted in 10% lymph glands (Fig. 3Q-Q’, T) with lamellocytes as compared to the control (Fig. 3O-O’, T). We then induced accelerated ageing in the *Chiz*-positive intermediate progenitor population and observed an increase in plasmatocyte differentiation upon Toll or Imd activation (Fig. S2D, E and P) whereas crystal cells showed an increase upon Imd activation and a suppression upon Toll pathway activation (Fig. S2I-J’ and Q) as compared to the control (Fig. S2A, F-F’, P-Q). Induction of decelerated ageing via Atg8 or Foxo over-expression had no effect on both plasmatocyte or crystal cell differentiation (Fig. S2B, C, G-H’, P and Q) as compared to the control (Fig. S2A, F-F’, P-Q). In terms of lamellocyte differentiation, induction of Toll or Imd pathway led to lamellocyte differentiation in the lymph glands (Fig. S2N-O’, R) whereas decelerated ageing had no induction of lamellocytes in the lymph glands (Fig. S2L-M’, R) as compared to their *wildtype* control (Fig. S2K-K’, R).

### Differentiated blood cell-specific modulation of ageing has cell autonomous effects on differentiation

Previous studies have suggested that differentiated hemocytes can reciprocally signal to the progenitors to regulate their numbers and thereby maintain homeostasis (Mondal et al., 2011). In order to understand if ageing modulation has dominant effects only on the niche and progenitors or can affect differentiated hemocytes cell-autonomously too we perturbed the ageing circuitry using *hmlΔGal4*, a differentiated hemocyte-specific *Gal4*. *hmlΔGal4* mediated Imd activation led to an increase in both plasmatocytes and crystal cells and resulted in LGs positive for lamellocytes (Fig. S3D-D’, K-K’, R-R’, F, M and T) whereas Toll activation led to a complete suppression of the plasmatocyte lineage, increase in crystal cells and a consequent 100% lamellocyte positivity in the lymph glands (Fig. S3E-E’, L-L’, S-S’, F-F’, M and T) as compared to the control (Fig. S3A-A’, H-H’, O-O’, F-F’, M and T). On the other hand, Atg8 over-expression using *hmlΔGal4* had no significant effect on all three differentiated hemocyte lineages (Fig. S3B-B’, I-I’, P-P’, G, N and T). Foxo over-expression resulted in an increase in plasmatocyte differentiation index, decrease in crystal cell numbers and a modest number of LGs (15%) showing lamellocyte positivity (Fig. S3C-C’, J-J’, Q-Q’, G, N and T) as compared to the *wildtype* control (Fig. S3A-A’, H-H’, O-O’, G, N and T).

### Systemic perturbation of ageing circuitry has an impact on lymph gland hematopoiesis

Organismal ageing could potentially impact multiple stem cell systems resulting in deregulated homeostasis. There is active systemic inter-organ communication and cross-talk between various organs. Lymph gland hemocytes have been shown to be responsive to external stimuli from peripheral organs (Benmimoun et al., 2012, Shim et al., 2012, Yang et al., 2015, Cho et al., 2018, Koranteng et al., 2022). Various reports point towards the involvement of fat body and muscles in determining overall organismal lifespan (Demontis & Perrimon, 2010, Bai et al., 2012, Owusu Ansah et al., 2013, Gáliková & Klepsatel., 2018, Guo et al., 2023). Here, we have used fat body and muscles as systemic sites for modulation of cellular ageing and tested the effects of the perturbation on blood cell homeostasis in the lymph gland. Fat body mediated induction of accelerated ageing by *pplGal4* results in an increase in plasmatocyte & crystal cell differentiation (Fig. S4D, E, I, J, P, R) and an induction of lamellocytes in the lymph glands as compared to the control. For lamellocytes, the activation of Toll drives lamellocytes in all the lymph glands whereas Imd activation in the fat body results in 18% of lymph glands having lamellocytes (Fig. S4N-O’, T) as compared to the control (Fig. S4A, F, K-K’, P, R and T). In the case of decelerated ageing, Atg8 over-expression in the fat body has no significant effect on hematopoiesis (Fig. S4B-L’, Q, S and T) whereas Foxo over-expression results in increased plasmatocytes (Fig. S4C, Q), no effect on crystal cells (Fig. S4H, S) and no induction of lamellocytes in the lymph glands (Fig. S4M-M’, T) as compared to the *wildtype* control (Fig. S4A-K’, P-T). Muscle- specific activation of both Toll or Imd leads to aberrant differentiation of plasmatocytes (Fig. 4D, E and P), an increase in crystal cells is observed upon Imd activation (Fig. 4I, Q) whereas no effect is seen upon Toll activation (Fig. 4J, Q) as compared to the control (Fig. 4A, F, P and Q). For lamellocytes, both Toll and Imd activation in the muscles triggers lamellocyte production (Fig. 4N-O’, R) in the lymph glands as compared to the control (Fig. 4K-K’, R). Interestingly, muscle- specific Foxo over-expression leads to a decrease in both plasmatocyte and crystal cell differentiation and no induction of lamellocytes in the lymph glands (Fig. 4C, H, M-M’, P-R). Atg8 over-expression does not cause any significant effect on the hemocyte differentiation (Fig. 4B, G, L-L’, P-R) as compared to the *wildtype* control (Fig. 4A, F, K-K’, P-R).

**Figure 4:**
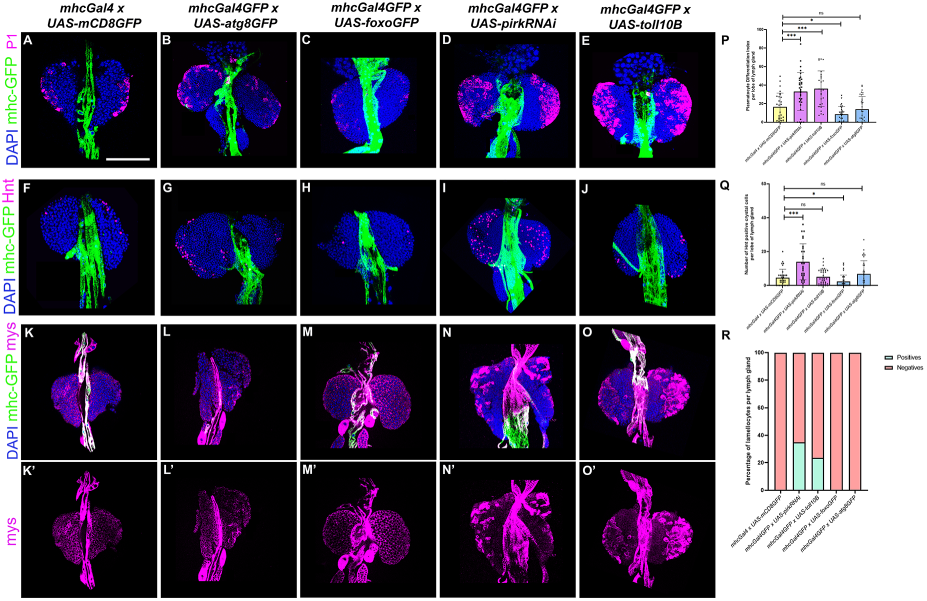
Muscle-specific systemic modulation of cellular ageing affects hemocyte differentiation in the LG. Plasmatocyte differentiation marked by P1 (magenta) or crystal cell differentiation marked by Hindsight (Hnt) (magenta) or lamellocyte differentiation marked by β-integrin-specific Myospheroid (Mys) (magenta) upon systemic muscle-specific myosin heavy chain (MHC, *mhcGal4*) mediated expression of *UAS-atg8GFP* (Fig. 4B, G, L), *UAS-foxoGFP* (Fig. 4C, H, M), *UAS-pirkRNAi* (Fig. 4D, I, N) or *UAS-toll10B* (Fig. 4E, J, O) as compared to *wildtype* control (Fig. 4A, F, K). Graphical representation of plasmatocyte differentiation index (Fig. 4P) or number of crystal cells (Fig. 4Q) or percentage of lamellocyte positive lymph glands (Fig. 4R) upon *mhcGal4GFP* mediated expression of *UAS-atg8GFP, UAS-foxoGFP, UAS-pirkRNAi* or *UAS-toll10B* as compared to *wildtype* control. For Plasmatocyte differentiation, *mhcGal4GFP x UAS-atg8GFP* (N = 16, n = 32), *mhcGal4 x UAS-foxoGFP* (N = 15, n = 30), *mhcGal4 x UAS-pirkRNAi* (N = 16, n = 32) and *mhcGal4 x UAS-toll10B* (N = 17, n = 34) were analyzed as compared to *mhcGal4 x UAS-GFP* (N = 14, n = 28). For crystal cell numbers, *mhcGal4 x UAS-atg8GFP* (N = 17, n = 34), *mhcGal4 x UAS-foxoGFP* (N = 16, n = 31), *mhcGal4 x UAS-pirkRNAi* (N = 15, n = 30) and *mhcGal4 x UAS-toll10B* (N = 17, n = 33) were analyzed as compared to *mhcGal4 x UAS-GFP* (N = 17, n = 33). For lamellocyte differentiation, *mhcGal4 x UAS-atg8GFP* (N = 19), *mhcGal4 x UAS-foxoGFP* (N = 17), *mhcGal4 x UAS-pirkRNAi* (N = 20) and *mhcGal4 x UAS-toll10B* (N = 17) were analyzed as compared to *mhcGal4 x UAS-GFP* (N = 16). N denotes the number of larvae & n denotes the number of individual primary lymph gland lobes analyzed per genotype. Individual data points in the graphs represent individual primary lobes of the Lymph gland. GFP expression (green) is driven by *mhcGal4* (Fig. 4A-O). Nuclei are stained with DAPI (Blue). Values are mean ± SD, and asterisks denote statistically significant differences (ns denotes not significant, **p < 0.01, *** p < 0.001, ****p < 0.0001). Student’s t-test with Welch’s correction was performed for the statistical analysis. Scale Bar: 50 µm (A-O’).

### Localized or systemic modulation of cellular ageing regulates DNA damage in the *Drosophila* lymph gland

One of the hallmarks of stem cell aging is oxidative stress-induced DNA damage in the form of Double stranded breaks (DSBs) which is marked by phosphorylated ***γ***-H2AX (Moehrle & Geiger., 2016, Carlos López-Otín et al., 2023). Previous studies have indicated age-related accumulation of DNA damage in intestinal stem cells (ISCs) of *Drosophila* (Park et al., 2012). Also, aged hematopoietic stem cells in vertebrates have been shown to accumulate DNA damage (Beerman et al., 2016). Here, we investigate the effect of modulating cellular aging in different cellular subsets of lymph gland and in distant organs like the fat body on the accumulation of DNA damage in the LG hemocytes. Induction of decelerated ageing by Foxo over-expression in the niche suppresses DNA damage accumulation whereas there is no significant effect when Foxo is over-expressed in the progenitors, whole lymph gland, differentiated hemocytes or in the fat body as compared to the respective controls. Activation of accelerated aging mediated by Imd pathway upregulation in the niche (Fig. S5C, D), medullary zone-specific progenitors (Fig. S5G,H), whole lymph gland (Fig. S5O, P) or fat body (Fig. S5S, T) exhibited an increase in the accumulation of DNA damage in LG hemocytes whereas differentiated hemocyte- specific Imd activation had no effect on DNA damage accumulation (Fig. S5K, L), as compared to respective *wildtype* controls (Fig. S5A-Q, D-T). Induction of decelerated ageing mediated by Foxo over-expression in the niche suppresses DNA damage accumulation (Fig. S5B, D) whereas its over-expression in the progenitors, whole lymph gland, differentiated hemocytes or in the fat body (Fig. S5F-R, H-T) has no significant effect as compared to the respective controls (Fig. S5A-Q, D-T). These results indicate that in the conditions of inflammatory insults including oxidative stress and elevated ROS levels induced by chronic activation of inflammatory pathways like Imd (Khor & Cai., 2020), there is severe accumulation of DNA damage (Shaposhnikov et al., 2015) whereas, in the case of Foxo over-expression which is known to upregulate cellular stress-response genes and DNA damage-response genes (Junger et al., 2003), very less DNA damage accumulation was observed, thereby further validating our genetic modalities of modulating cellular aging.

### Localized or systemic induction of proteotoxic stress impairs blood cell homeostasis in Drosophila

In order to support our findings on inducing accelerated ageing by Toll or Imd activation that induces inflammaging, we used an alternative approach of inducing accelerated aging by loss of proteostasis and generation of proteotoxic stress due to accumulation of misfolded protein aggregates. Perturbation of proteostasis and accumulation of protein aggregates is a key hallmark of cellular aging (Carlos López-Otín et al., 2023). We chose Huntington’s model as a medium to genetically induce proteotoxic stress based on previous studies that indicated that mutated huntingtin (HTT) protein in the Huntington’s autosomal dominant neurodegenerative disease is over-expressed and misfolds and is prone to aggregation and accumulation, thereby causing impairment in proteostasis which is otherwise involved under normal conditions in protein folding and clearance and degradation of HTT aggregates (Koyuncu, 2017) and the ability to maintain proteostasis in the background of HTT aggregate overload in Huntington’s disease declines with aging in induced pluripotent stem cells (iPSCs) (Mattis et al., 2013).

Here, we expressed mutated version of the Huntingtin (HTT) protein (causing Huntington’s disease in mammals) in the niche or hematopoietic progenitors to ascertain the effect of proteotoxic stress on blood cell homeostasis. Upon genetic induction of HTT-RFP expression in the niche, there was a significant reduction in the niche size (Fig. 5B-10B’, 10C) in a cell-autonomous manner with a concomitant increase in plasmatocyte differentiation (Fig. 5E, F), increase in crystal cell numbers (Fig. 5H, I) and 95% lymph glands positive for lamellocytes (Fig. 5K-10K’, L) as compared to the *wildtype* control (Fig. 5A-J’, C-L). Similarly, induction of HTT aggregates in *tepIV*-positive core-progenitor pool exhibited a non-autonomous decrease in niche size (Fig. 5N, O), cell-autonomous reduction in progenitor pool (Fig. 5Q, R) and an increase in both plasmatocyte (Fig. 5T, U) and crystal cell differentiation (Fig. 5W, X). Induction of HTT expression in progenitors also results in 50% lymph glands positive for lamellocytes (Fig. 5Z-Z”) as compared to *wildtype* control (Fig. 5M-Y’, O-Z”). These observations from genetic induction of HTT aggregates in different subsets of lymph gland promoted accelerated aging mediated by proteotoxic stress and supports our data on inflammaging thereby leading to perturbation of blood cell homeostasis in *Drosophila*. Systemic induction of proteotoxic stress by expression of HTT-RFP results in a decrease in niche cell numbers (Fig. S6B-C), increase in both plasmatocyte (Fig. S6E-F) and crystal cell differentiation (Fig. S6H-I) and induces lamellocytes in 100% of the lymph glands analyzed (Fig. S6K-K’, L) as compared to the *wildtype* control (Fig. S6A-J’, C-L).

**Figure 5:**
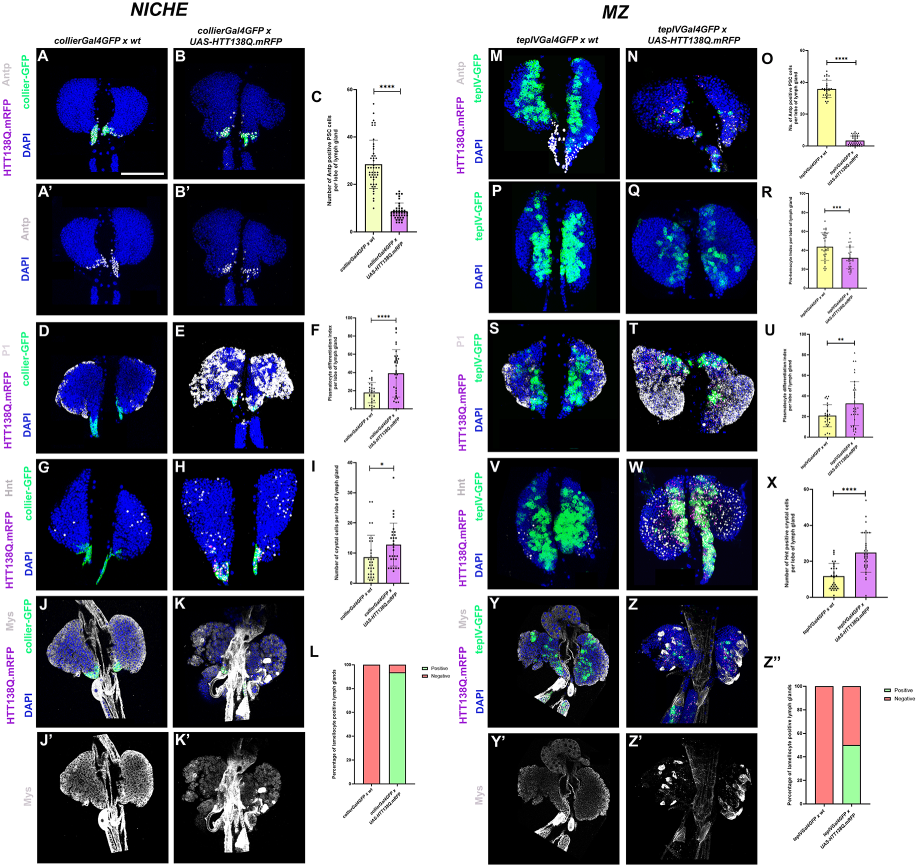
Genetic induction of proteotoxic stress in the stem cell niche or hematopoietic progenitors results in aberrant hematopoiesis. Posterior Signalling Centre (PSC) niche cell population marked by Antennapedia (white, Fig. 5B-B’, N) or Plasmatocyte differentiation marked by P1 (white, Fig. 5E, T) or Crystal cell marked by Hindsight (Hnt in white, Fig. 5H, W) or lamellocyte differentiation marked by β-integrin specific Myospheroid (Mys in white, Fig. 5K-K’, Z-Z’) upon genetic induction of proteotoxic stress by expression of *HTT138Q.mRFP* in either PSC niche-specific (using *collierGal4*) or Medullary zone core progenitor-specific (using *tepIVGal4*) manner as compared to respective wild type control (Fig. Niche – 5A-A’, D, G, J-J’; MZ – 5M, S, V, Y-Y’). tep positive hematopoietic progenitors (Green) index upon *tepIVGal4* mediated expression of *HTT138Q.mRFP* as compared to *wild type* control (Fig. 5P-R). *HTT138Q.mRFP* is marked in magenta and nuclei are stained with DAPI (Blue). Graphical representation of PSC niche numbers (Fig. 5C, O), plasmatocyte differentiation index (Fig. 5F, U), Crystal cell numbers (Fig. 5I, X) and percentage of lamellocyte positive lymph glands (5L, Z”) upon genetic induction of proteotoxic stress in either PSC niche or pro-hemocyte population as compared to their respective wildtype control (Fig. 5C, F, I, L, O, R, U, X, Z”). For genetic induction of proteotoxic stress in either PSC niche or core-progenitor population, the PSC niche numbers (Niche: *collierGal4 x wt* – N = 24, n = 31; *collierGal4 x UAS-HTT138Q.mRFP* - N = 20, n = 40, **MZ**: *tepIVGal4 x wt* – N = 24, n = 48; *tepIVGal4 x UAS-HTT138Q.mRFP* – N = 19, n = 38) or plasmatocyte differentiation (Niche: *collierGal4 x wt* – N = 14, n = 28; *collierGal4 x UAS-HTT38Q.mRFP* - N = 18, n = 36, **MZ**: *tepIVGal4 x wt* – N = 16, n = 32; *tepIVGal4 x UAS-HTT38Q.mRFP* – N = 18, n = 36) or Crystal cell numbers (Niche: *collierGal4 x wt* – N = 15, n = 30; *collierGal4 x UAS-HTT138Q.mRF*P – N = 15, n = 29, **MZ**: *tepIVGal4 x wt* – N = 16, n = 31; *tepIVGal4 x UAS-HTT*138Q.m*RFP* – N = 17, n = 33) or lamellocyte differentiation (Niche: *collierGal4 x wt* – N = 15; *collierGal4 x UAS-HTT*138Q.m*RFP* - N = 16, **MZ**: *tepIVGal4 x wt* – N = 14.5; *tepIVGal4 x UAS-HTT*138Q.m*RFP* – N = 20) were analyzed per genotype. For analysis of core-progenitor index upon genetic induction of proteotoxic stress in core-progenitor population, *tepIVGal4 x wt* – N = 15, n = 30 and *tepIVGal4 x UAS-HTT*138Q.m*RFP* – N = 15, n = 30 were utilised. N denotes the number of larvae & n denotes the number of individual primary lymph gland lobes analysed per genotype. GFP expression (green) is either driven by *collierGal4* (Fig. 5A- K) or *tepIVGal4* (Fig. 5M - Z). Nuclei are marked with DAPI (Blue). Individual data points in the graphs represent individual primary lobes of the Lymph Gland. Values are mean ± SD, and asterisks denote statistically significant differences (ns denotes not significant, **p < 0.01, *** p < 0.001, ****p < 0.0001). Student’s t-test with Welch’s correction was performed for the statistical analysis. Scale Bar: 50 µm (A-K’, M-Z’).

### Modulation of ageing by chemical-based intervention regulates blood cell homeostasis in the lymph gland

In order to supplement our findings on genetic modulation of ageing we also used chemical intervention as an approach to modulate ageing. Here, we have used Rapamycin for decelerating aging and Bortezomib treatment for accelerating aging at an organismal level and have investigated its effect on blood cell homeostasis in the LG. Previous studies have shown that Bortezomib acts as an inhibitor of the ubiquitin-mediated proteasomal pathway (Chen et al., 2011), thereby resulting in increased protein instability, redox imbalance and accelerated aging (Manola et al., 2019). Additionally, Bortezomib was also shown to induce cellular senescence by stimulating telomere shortening in non-small cell lung cancer (NSCLC) cells (Yin et al., 2022). Bortezomib treatment in *wildtype* control larvae with green fluorescence marking the *tepIV*-positive core progenitors exhibited significant increase in plasmatocyte (Fig. 6A’-A”) and crystal cell differentiation (Fig. 6B’-B”) when compared to its vehicle control (Fig. 6A, A’’, B, B”) indicating that there is perturbation of LG homeostasis upon inhibition of proteostasis. However, rapamycin treatment in *wildtype* larvae with green fluorescence marking the *tepIV*-positive core progenitors showed a reduction in plasmatocyte differentiation (Fig. 6C’-C”), whereas there was no effect observed on crystal cell differentiation (Fig. 6D’- D”) as compared to its vehicle control (Fig. 6C, C”, D, D”). Previous literature has shown that Rapamycin inhibits mTORC1 (mammalian Target of Rapamycin Complex1) which is generally upregulated in amino acid-rich conditions and promotes protein synthesis and inhibits autophagy (Wu et al., 2013). Inhibition of mTORC1 kinase by Rapamycin leads to upregulation of autophagy, increased resistance to starvation and lifespan extension in *Drosophila* (Bjedov et al., 2010). After testing the effect of Rapamycin on *wildtype* control larvae, we wanted to test if Rapamycin could rescue the blood cell differentiation defects in larvae with Toll or Imd activation in the progenitors. Rapamycin administration to larvae having Imd activation in the *tepIV*-positive core progenitors could rescue plasmatocyte (Fig. 6E’-E”) and crystal cell differentiation significantly (Fig. 6G’-G”) as compared to their control (Fig. 6E, G, E’’ and G’’). However, rapamycin did not exhibit any rescue of differentiation for lamellocytes (Fig. 6I’- I”) when compared to its vehicle control (Fig. 6I, I”). Similarly, administration of rapamycin in larvae with Toll pathway over-activation in *tepIV*-positive core progenitors was unable to rescue the Toll-mediated differentiation of all three hemocyte lineages (Fig. 6F’- J”) as compared to its vehicle control (Fig. 6F-J’, F’’-J’’).

**Figure 6:**
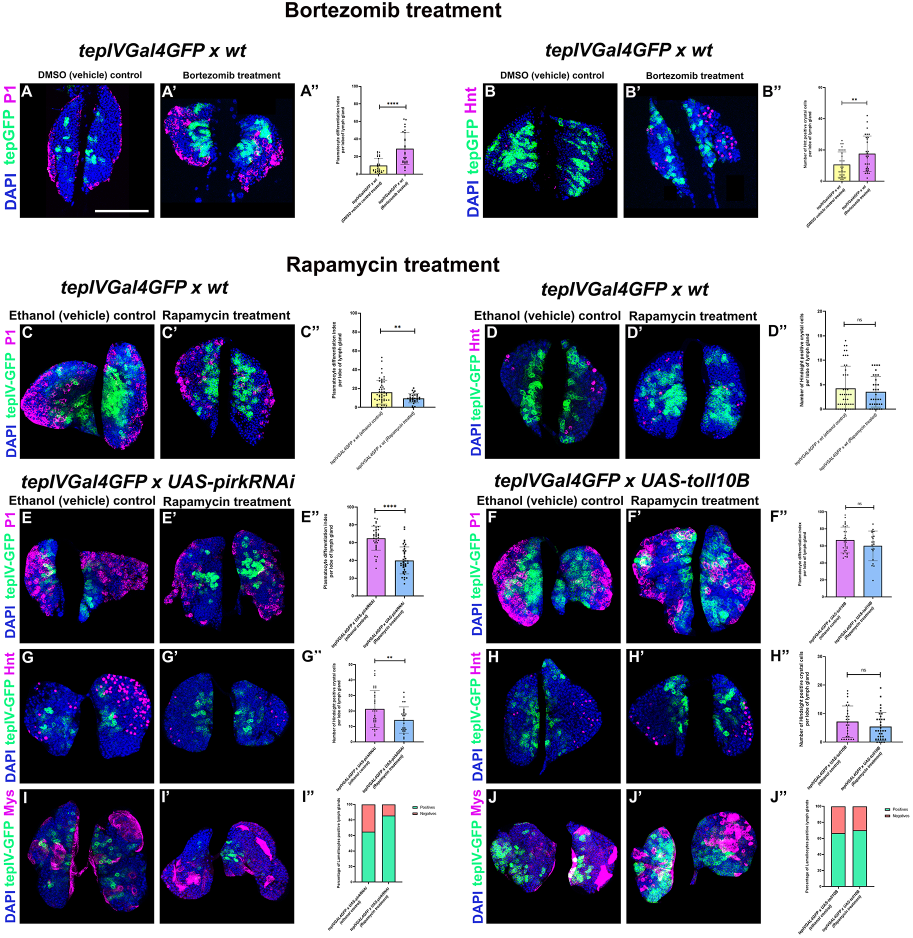
Chemical modulation of cellular ageing affects blood cell homeostasis in *Drosophila*. Plasmatocyte differentiation marked by P1 (magenta) or crystal cell differentiation marked by Hindsight (Hnt, magenta) in *tepIVGal4 x wt* larvae upon treatment with Bortezomib (Fig. 6A’- B’) as compared with DMSO (vehicle) control (Fig. 6A - B). Graphical representation of Plasmatocyte differentiation index (Fig. 6A”) or number of crystal cells (Fig. 6B”) upon Bortezomib treatment as compared with DMSO (vehicle) control. For plasmatocyte differentiation upon bortezomib treatment, *tepIVGal4 x wt* (N = 15, n = 29) were analyzed as compared to DMSO (vehicle) control (N = 13, n = 26). Plasmatocyte differentiation marked by P1 (magenta) or crystal cell differentiation marked by Hindsight (Hnt, magenta) in *tepIVGal4 x wt* (Fig. 6C’, D’) or *tepIVGal4 x UAS-pirkRNAi* (Fig. 6E’, G’) or *tepIVGal4 x UAS-toll10B* (Fig. 6F’, H’) larvae upon treatment with Rapamycin as compared with ethanol (vehicle) control (Fig. 6C-D, E - H). Percentage of Lamellocyte positive lymph glands in *tepIVGal4 x UAS-pirkRNAi* (Fig. 6I’) or *tepIVGal4 x UAS-toll10B* (Fig. 6J’) upon treatment with Rapamycin as compared with ethanol (vehicle) control (Fig. 6I-J). Graphical representation of plasmatocyte differentiation index or number of crystal cells upon Rapamycin treatment in *tepIVGal4 x wt* (Fig. 6C”- D”) or *tepIVGal4 x UAS-pirkRNAi* (Fig. 6E”, G”) or *tepIVGal4 x UAS-toll10B* (Fig. 6F”, 6H”) larvae or percentage of lamellocyte positive lymph glands in *tepIVGal4 x UAS-pirkRNAi* (Fig. 6I”) or *tepIVGal4 x UAS-toll10B* (Fig. 6J”) larvae upon treatment with Rapamycin as compared with ethanol (vehicle) control. For plasmatocyte differentiation upon Rapamycin treatment, *tepIVGal4 x wt* (N = 15, n = 30), *tepIVGal4 x UAS-pirkRNAi* (N = 19, n = 38), *tepIVGal4 x UAS-toll10B* (N = 10, n = 20) were analyzed as compared to respective ethanol (vehicle) controls: *tepIVGal4 x wt* (N = 25, n = 50), *tepIVGal4 x UAS-pirkRNAi* (N = 18, n = 36), *tepIVGal4 x UAS-toll10B* (N = 17, n = 34). For crystal cell differentiation, *tepIVGal4 x wt* (N = 20, n = 39), *tepIVGal4 x UAS-pirkRNAi* (N = 13, n = 26), *tepIVGal4 x UAS-toll10B* (N = 18, n = 35) were analyzed as compared to respective ethanol (vehicle) controls: *tepIVGal4 x wt* (N = 24, n = 48), *tepIVGal4 x UAS-pirkRNAi* (N = 17, n = 34), *tepIVGal4 x UAS-toll10B* (N = 14, n = 28). For Lamellocyte differentiation, *tepIVGal4 x UAS-pirkRNAi* (N = 15) and *tepIVGal4 x UAS-toll10B* (N = 15) were analyzed as compared to respective ethanol (vehicle) controls: *tepIVGal4 x UAS-pirkRNAi* (N = 15) and *tepIVGal4 x UAS-toll10B* (N = 15). N denotes the number of larvae & n denotes the number of individual primary lymph gland lobes analysed per genotype. Individual data points in the graphs represent individual primary lobes of the lymph Gland. GFP (green) is driven by *tepIVGal4* which marks pro-hemocytes (Fig. 6A - J, Fig. 6A’ - J’). Nuclei are stained with DAPI (Blue). Values are mean ± SD, and asterisks denote statistically significant differences (ns denotes not significant, **p < 0.01, *** p < 0.001, ****p < 0.0001). Student’s t-test with Welch’s correction was performed for the statistical analysis. Scale Bar: 50 µm (A-J’).

### Genetic modulation of cellular ageing regulates Integrated Stress Response (ISR) signalling in the lymph gland

The Integrated stress response pathway (ISR) is an important and highly conserved cellular signaling pathway that integrates various signaling pathways that counter various stressors and provides a combined and centralized cellular response that facilitates the organism to adapt to various environmental stresses and promotes maintenance of cellular bioenergetics until the stress is resolved, thereby aiding in survival (Pakos-Zebrucka et al., 2016, Kalinin et al., 2023). Previous studies have shown that as age advances, the ISR pathway is upregulated and Gcn2, eIF2ɑ & Crc-RA (a splice variant and isoform of *crc* gene which is a *Drosophila* ortholog of *atf4*) (Vasudevan et al., 2022) phosphorylation was increased in the brain of old mice. It was also reported that ISR activation facilitated lifespan extension in *Drosophila* and *C. elegans* (Derisbourg et al., 2022). Interestingly, it has also been found that prolonged ISR activation due to conditions of excessive and persistent stress contributed to cell senescence or programmed cell death (Kalinin et al., 2023). Previous studies have also reported the activation of ISR upon enhanced inflammation and cytokine secretion in mice and ISR activation and phosphorylation of eIF2ɑ was shown to inhibit the translation of cytokine mRNAs in the tissue-resident memory T cells (Trm) of mice (Asada et al., 2024).

In order to investigate if the stress caused by chronic activation of Toll or Imd that abrogates autophagy, protein turnover and elevates ROS levels results in ISR activation or not, we probed for p-eIF2α levels in the lymph glands with Toll or Imd activation or upon induction of decelerated ageing in the hematopoietic progenitors. Our results indicate that the downstream readout of the ISR pathway i.e. p-eIF2ɑ levels measured by quantitating the mean fluorescence intensity in the *tepIV*-positive core progenitors was significantly increased upon Toll or Imd activation (Fig. 7B-C’, G) whereas over-expression of Atg8 (Fig. 7D-D’, G) or Foxo (Fig. 7E-E’, G) showed no significant difference as compared to the *wildtype* control (Fig. 7A-A’, G). In addition to this, we also profiled for *gcn2* transcript levels from the hemolymph isolated from larvae where either accelerated or decelerated ageing was induced in all the blood cells using *hmlGal4*, a pan hemocyte *Gal4*. *gcn2* mRNA levels were upregulated in the inflammaging scenario i.e. during the Imd or Toll pathway activation, while Atg8 or Foxo over-expression showed suppression in the Gcn2 levels as compared to *wildtype* control (Fig. 7F). This suggests that the upstream ISR activator, Gcn2 and downstream ISR key effector, p-eIF2ɑ are downregulated or inactive in the decelerated aging scenario as the cells were already compensating and countering stresses via ectopic over-expression of Atg8 or Foxo whereas ISR is much needed in an inflammaging scenario to possibly sense and counter various stresses and promote cell survival.

**Figure 7:**
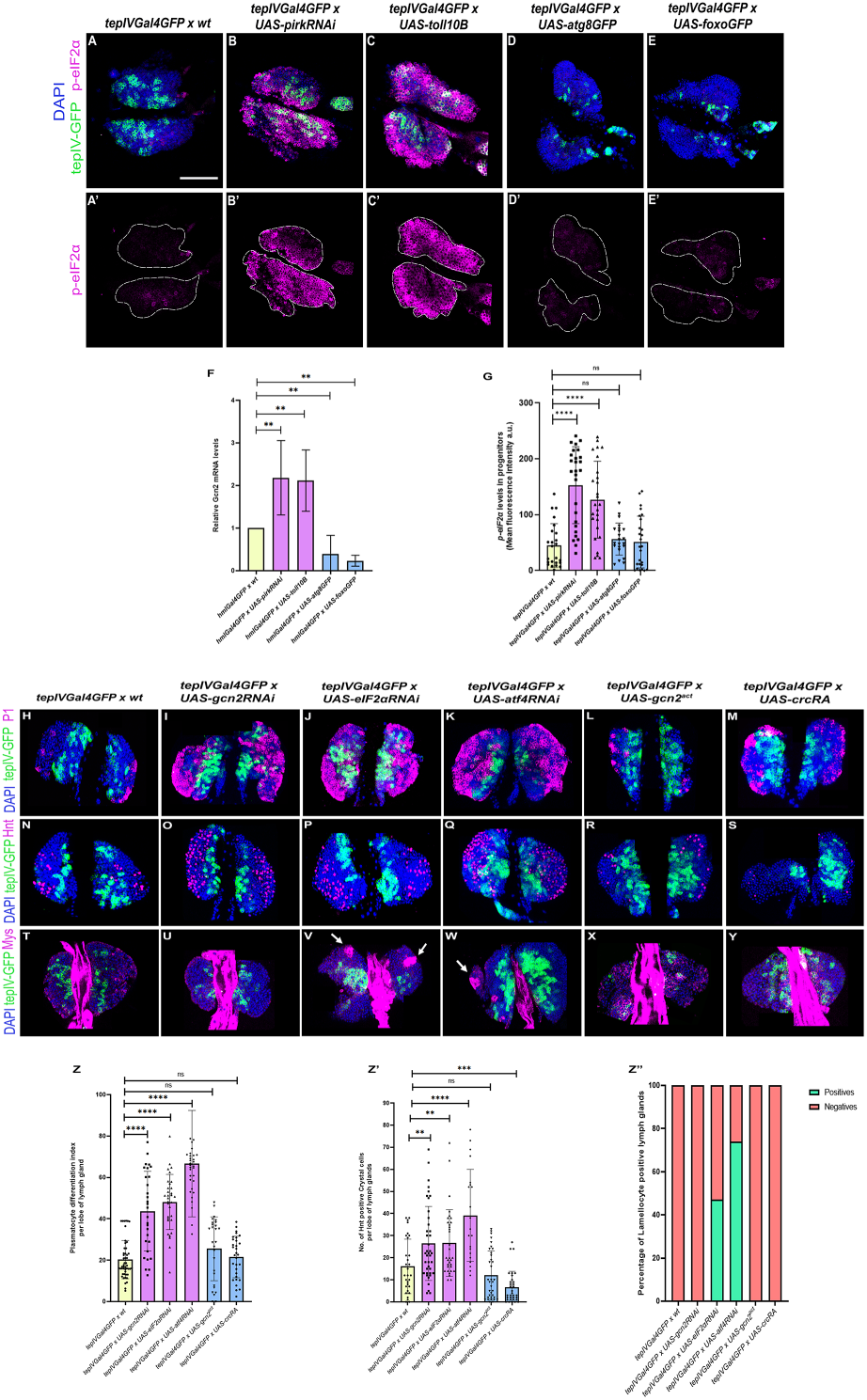
Hematopoietic progenitor- specific induction of accelerated ageing triggers Integrated Stress Response (ISR) pathway that regulates progenitor differentiation in the LG. Mean fluorescence intensity of p-eIF2α levels in the core progenitors marked by p-eIF2α antibody (magenta) upon induction of accelerated aging using (*UAS-pirkRNAi* or *UAS- toll10B*) or decelerated aging (*UAS-atg8* or *UAS-foxo*) as compared to *wildtype* control (Fig. 7A-E, A’-E’, G). Graphical representation of p-eIF2α fluorescence intensity levels upon *tepIVGal4* mediated induction of accelerated aging (*UAS-pirkRNAi* or *UAS-toll10B*) or decelerated aging (*UAS-atg8* or *UAS-foxo*) as compared to *wildtype* control (Fig. 7G). Relative *gcn2* mRNA transcript levels determined by qRT-PCR upon pan-hemocyte-specific (using *hmlGal4*) expression of *UAS-atg8GFP*, *UAS-foxoGFP*, *UAS-pirkRNAi* or *UAS-toll10B* as compared to *wildtype* control (Fig. 7F). The statistical analysis for qRT-PCR was performed using One-way ANOVA (Dunnett test) for comparison of all test genotypes with *wildtype* control. Plasmatocyte differentiation marked by P1 (magenta) or crystal cell differentiation marked by Hindsight (Hnt, magenta) or lamellocyte differentiation marked by β-integrin- specific Myospheroid (Mys, magenta) upon core progenitor-specific (using *tepIVGal4*) knockdown (*gcn2RNAi, atf4RNAi or eIF2αRNAi)* and over-expression (*gcn2^act^or crcRA*) of ISR components (Fig. 7I-Y, Z-Z’’) as compared to *wildtype* control (Fig. 7H, N, T, Z-Z’’). Graphical representation of plasmatocyte differentiation index (Fig. 7Z) or number of crystal cells (Fig. 7Z’) or percentage of lamellocyte positive lymph glands (Fig. 7Z’’) upon *tepIVGal4* mediated knockdown or over-expression of ISR components as compared to *wildtype* control. For p-eIF2α levels, *tepIVGal4 x UAS-pirkRNAi* (N = 14, n = 27), *tepIVGal4 x UAS-toll10B* (N = 13, n = 26), *tepIVGal4 x UAS-atg8GFP* (N = 11, n = 22) and *tepIVGal4 x UAS-foxoGFP* (N = 13, n = 26) were analyzed as compared to *tepIVGal4 x wt* (N = 12, n = 24) *as wildtype* control. For plasmatocyte differentiation, *tepIVGal4 x UAS-gcn2RNAi* (N = 17, n = 34), *tepIVGal4 x UAS-eIF2αRNAi* (N = 16, n = 32), *tepIVGal4 x UAS-atf4RNAi* (N = 16, n = 32), *tepIVGal4 x UAS-gcn2^act^* (N = 15, n = 30), *tepIVGal4 x UAS-crcRA* (N = 18, n = 36) were analyzed as compared to *wildtype* control, *tepIVGal4 x wt* (N = 25, n = 49). For crystal cell differentiation, *tepIVGal4 x UAS-gcn2RNAi* (N = 20, n = 40), *tepIVGal4 x UAS-eIF2αRNAi* (N = 17, n = 33), *tepIVGal4 x UAS-atf4RNAi* (N = 13, n = 25), *tepIVGal4 x UAS-gcn2^act^* (N = 18, n = 35), *tepIVGal4 x UAS-crcRA* (N = 17, n = 33) were analyzed as compared to *wildtype* control, *tepIVGal4 x wt* (N = 18, n = 36). For lamellocyte differentiation, *tepIVGal4 x UAS-gcn2RNAi* (N = 15, n = 30), *tepIVGal4 x UAS-eIF2αRNAi* (N= 15, n = 30), *tepIVGal4 x UAS-atf4RNAi* (N = 15, n = 30), *tepIVGal4 x UAS-gcn2^act^* (N = 15, n = 30), *tepIVGal4 x UAS-crcRA* (N = 15, n = 30) were analyzed as compared to *wildtype* control, *tepIVGal4 x wt* (N = 15, n = 30). N denotes the number of larvae & n denotes the number of individual primary lymph gland lobes analyzed per genotype. Individual data points in the graphs represent individual primary lobes of the Lymph gland. GFP expression (green) is driven by *tepIVGal4* (Fig. 7A-Y). Nuclei are stained with DAPI (Blue). Values are mean ± SD, and asterisks denote statistically significant differences (ns denotes not significant, **p < 0.01, *** p < 0.001, ****p < 0.0001). Student’s t-test with Welch’s correction was performed for the statistical analysis. Scale Bar: 50 µm (A-Y).

### Progenitor-specific genetic perturbation of ISR pathway components regulates LG hematopoiesis

Since ISR pathway components and downstream effectors are induced in an accelerated ageing scenario, we wanted to further test if depletion or over-expression of ISR pathway components in the hematopoietic progenitors has any impact on LG hematopoiesis. Depletion of Gcn2, eIF2α or Atf4 led to an increase in both plasmatocyte and crystal cell differentiation (Fig. 7I, J, K, O, P, Q, Z-Z’) whereas over-expression of *gcn2^act^* has no effect on plasmatocyte or crystal cell differentiation (Fig.7L, R, Z-Z’), over-expression of *crcRA* has no effect on plasmatocyte differentiation but results in a decrease in crystal cell differentiation (Fig.7M, S, Z-Z’) as compared to the *wildtype* control (Fig.7H, N, Z-Z’). In terms of lamellocyte differentiation, depletion of eIF2α or Atf4 leads to lymph glands positive for lamellocytes (Fig. 7V, W, Z’’) whereas depletion of Gcn2 or over-expression of *gcn2^act^* or *crc-RA* (Fig. 7U, X, Y, Z’’) does not induce lamellocytes in the lymph glands as compared to the *wildtype* control (Fig. 7T, Z’’).

### Mutants of the ISR pathway components show disruption in blood cell homeostasis

In the previous results, we observed that depletion of ISR pathway components led to aberrant hematopoiesis. Following these observations, we set out to look at the lymph glands of whole animal mutants of *gcn2* and *crc* (encoding *Drosophila* Atf4). *gcn2^-/-^* homozygous null mutant and *crc^1^/+* (cryptocephal) heterozygous hypomorphic allele that have been previously reported were used and their lymph gland phenotypes were analyzed (Vasudevan et al., 2022). *gcn2^-/-^* homozygous null mutant lymph glands showed an increase in plasmatocyte (Fig. S7B-C) as well as crystal cell differentiation (Fig. S7E-F) and a presence of lamellocytes in 60% of the lymph glands (Fig. S7H-I) as compared to *gcn2^wt^* rescue; *gcn2^-/-^* as the control (Fig. S7A, D, G, C, F and I). Heterozygotes of *crc^1/+^* hypomorphic allele displayed a significant increase in plasmatocyte (Fig. S7K-L) as well as crystal cell differentiation (Fig. S7N-O) along with 80% lymph glands showing the presence of lamellocytes (Fig. S7Q-R) as compared to *wildtype* control (Fig. S7J, M, P, L, O and R).

### Hematopoiesis in the lymph gland is sensitive to chemical modulators that regulate the ISR pathway

Since our data on the genetic perturbation of ISR showed an effect on LG hematopoiesis we wanted to validate these findings using known pharmacological modulators of the ISR pathway. We have used the small-molecule ISRIB (ISR Inhibitor) & Histidinol (ISR activator) for our experiments. ISRIB desensitizes eIF2B to the inhibitory effect of p-eIF2ɑ and turns off ISR pathway (Zyryanova et al., 2021) whereas Histidinol which is an analog of Histidine activates the ISR pathway by mimicking amino acid starvation stress (Taniuchi et al., 2016). Upon ISRIB administration to *wildtype* larvae with hematopoietic progenitors marked with *tepIV-GFP*, there was a significant increase in progenitor differentiation into plasmatocytes (Fig. S8B-C), crystal cells (Fig. S8E-F) and about 80% lymph glands positive for lamellocyte production (Fig. S8H-I) when compared to its vehicle control (Fig. S8A, D, G-G’, C, F and I).

Now, since we observed that there is ISR activation upon inducing accelerated ageing in the hematopoietic progenitors we hypothesized that larvae where the progenitors encounter inflammaging conditions induced via Toll or Imd upregulation when subjected to sustained activation of ISR pathway can potentially counter the stress conditions. Such aged progenitors when exposed to sustained ISR pathway activation could possibly counter the stress and maintain blood cell homeostasis. In order to test this hypothesis, we treated larvae bearing Toll or Imd activation in the hematopoietic progenitors with Histidinol, an ISR pathway activator. Histidinol treatment to larvae with Toll activation in the progenitors leads to a rescue of plasmatocyte differentiation phenotype (Fig. S8K-L) whereas crystal cell numbers show no difference (Fig. S8N-O) as compared to the vehicle treatment to larvae with progenitor-specific Toll activation (Fig. S8J, M, L, O). There was also a moderate decrease in the percentage of lymph glands positive for lamellocytes upon Histidinol treatment (Fig. S8Q-Q’, R) as compared to the vehicle control (Fig. S8P-P’, R). Similarly, we treated the larvae with Imd activation in progenitors with Histidinol which resulted in a rescue in both plasmatocyte (Fig. S8T, U) as well as crystal cell differentiation (Fig. S8W, X). In this case, the percentage of lymph glands positive for lamellocytes also decreased (Fig. S8Z-Z’’) upon Histidinol treatment as compared to the vehicle control (Fig. S8Y-Y’, Z’’) where larvae with Imd activation in progenitors are treated with the vehicle alone.

### Ectopic hyperactivation of the ISR pathway in an accelerated ageing scenario restores blood cell homeostasis in the lymph gland

Our pharmacological intervention data suggests that sustained ISR activation in accelerated ageing background can rescue blood cell homeostasis. In order to validate if this holds true upon genetic activation of ISR pathway in the accelerated ageing background, we over-expressed the activated form of Gcn2 or CrcRA (a splice variant and isoform of *crc* gene that encodes *Drosophila* Atf4) (Vasudevan et al., 2022) in the progenitors where Imd is activated to check if the hematopoiesis parameters are altered. We observe that over-expression of *gcn2^act^*or *crcRA* in progenitors where Imd is over-activated leads to a rescue of plasmatocyte (Fig. 8B, C, D) and crystal cell differentiation (Fig. 8F, G, H) as compared to the control where Imd is over-activated in progenitors (Fig. 8A, D, E, H). We do not find any significant rescue of the lamellocyte phenotype upon over-expression of *gcn2^act^*or *crcRA* in the Imd over-activated genetic background in the progenitors (Fig. 8J-K’, L) as compared to the respective control (Fig.8I-I’, L). These results indicate that both chemical intervention and genetic approaches to activate ISR are capable of restoring blood cell homeostasis in lymph glands where inflammaging is activated in hematopoietic progenitors. In addition to the hematopoietic parameters, we also assessed if quality control mechanisms like autophagy that are perturbed in an ageing scenario are restored upon ISR activation. As per our previous results we had shown that autophagy is affected in an accelerated ageing scenario (Fig 1). Here, we over-expressed *gcn2^act^*or *crcRA* in the progenitors where Imd is over-activated and scored for p62, an adapter molecule that gets degraded during autophagy and the effector, Atg8. Our analysis indicates that p62 positive puncta per cell are lower upon *gcn2^act^* or *crcRA* over-expression in the Imd over-activated background in the progenitors (Fig.8N-O’, S) as compared to the control (Fig. 8M-M’, S) whereas Atg8 shows an opposite trend as compared to the respective control which shows that autophagy levels are restored upon ISR activation in the inflammaging scenario (Fig. 8P-R’, T). Our findings demonstrate that modulation of cellular ageing i.e. both accelerated or decelerated ageing have distinct effects on LG hematopoiesis and Integrated Stress Response (ISR) pathway is crucial for responding to stress inflicted by inflammaging in an accelerated ageing scenario. Sustained ISR activation is capable of countering the stress and restoring blood cell homeostasis in the LG (Fig. 8U).

**Figure 8:**
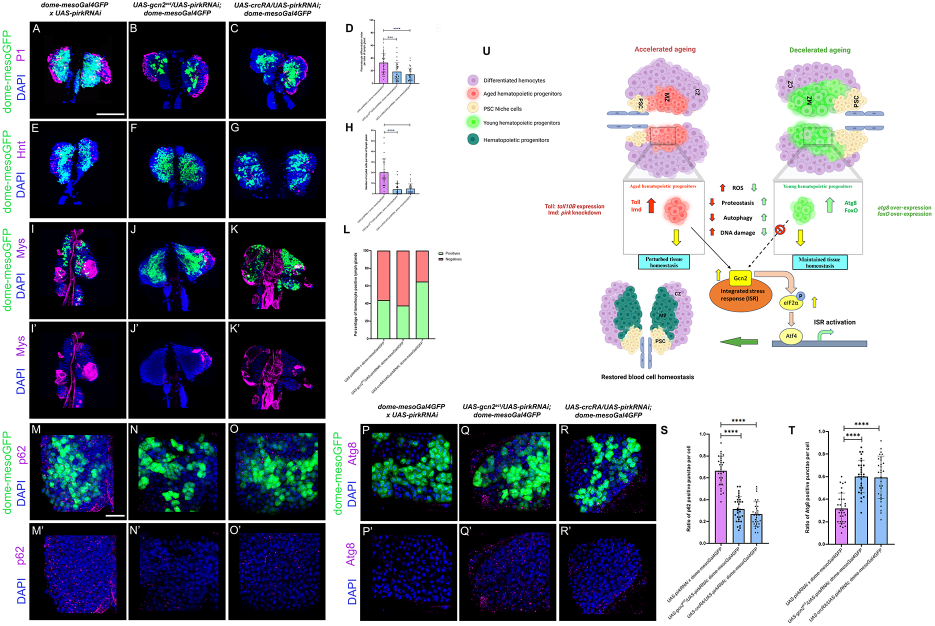
Sustained activation of ISR pathway in an accelerated ageing scenario in hematopoietic progenitors is capable of restoring blood cell homeostasis. Plasmatocyte differentiation marked by P1 (magenta) or crystal cell differentiation marked by Hindsight (Hnt, magenta) or lamellocyte differentiation marked by β-integrin-specific Myospheroid (Mys, magenta) or evaluation of p62 or Atg8 positive punctae per cell marked by p62 or Atg8 antibody (magenta) upon distal progenitor-specific (using *dome-mesoGal4*) hyperactivation of components of ISR pathway viz. *gcn2^act^*& *crcRA* in an IMD upregulated background (via *pirkRNAi*) as compared to *wildtype* control (Fig. 8A-K’, M-O’, P-R’). Graphical representation of plasmatocyte differentiation index or number of crystal cells or percentage of lamellocyte positive lymph glands or ratio of p62 or Atg8 positive punctae per cell upon *dome-mesoGal4* mediated over-expression of *gcn2^act^*& *crcRA* in a *pirk* knockdown background as compared to *wildtype* control (Fig.8D,H,L,S,T). *UAS-gcn2^act^/UAS- pirkRNAi; dome-mesoGal4 & UAS-crcRA/UAS-pirkRNAi; dome-mesoGal4* were analyzed for plasmatocyte differentiation (*gcn2^act^* : N = 15, n = 30; *crcRA:* N = 19, n = 37) or crystal cell numbers (*gcn2^act^* : N = 17, n = 33; *crcRA:* N = 14, n = 28) or lamellocyte differentiation (*gcn2^act^* : N = 16, n = 32; *crcRA:* N = 15, n = 30) or p62 levels (*gcn2^act^* : N = 11; *crcRA:* N = 10) or Atg8 levels per cell (*gcn2^act^*: N = 10; *crcRA:* N = 12) as compared to *wildtype* control (plasmatocytes: N = 19, n = 38; crystal cells: N = 14, n = 28; lamellocytes: N = 15, n = 30; p62: N = 12; Atg8: N = 10). N denotes the number of larvae & n denotes the number of individual primary lymph gland lobes analyzed per genotype. Individual data points in the graphs represent individual primary lobes of the Lymph gland. GFP expression (green) is driven by *dome-mesoGal4* (Fig. 8A-K’, M-O’, P-R’). Nuclei are stained with DAPI (Blue). Values are mean ± SD, and asterisks denote statistically significant differences (ns denotes not significant, **p<0.01, *** p<0.001, ****p<0.0001). Student’s t-test with Welch’s correction was performed for the statistical analysis. Our working model summarizing how restoration of blood cell homeostasis in the lymph gland occurs upon ectopic ISR hyperactivation in the hematopoietic progenitors in an accelerated ageing genetic background (NF-κB (Imd) upregulated) (Fig. 8U). Scale Bar: 50 µm (A-K’), 30 µm (M-R’).

## Discussion

One of the key hallmarks of ageing is stem cell exhaustion where the ability of self renewal of stem cells is compromised leading to aberrant differentiation and lack of maintenance of the stem cell pool that would otherwise be readily available in adverse conditions (Carlos Lopez-Otin., 2023, Yi et al., 2020). Here, in this study leveraging the strength of *Drosophila* genetics and using the hematopoietic system as our model we have done a systematic characterization of how genetic modulation of the ageing circuitry in various cellular subsets of the larval lymph gland (LG) can impact overall organ homeostasis. We not only modulate the ageing circuitry in a localized manner but also investigate how systemic modulation of ageing can regulate hematopoiesis via inter-organ communication. We use two genetic approaches – accelerated and decelerated ageing to answer these questions using the inflammaging approach for accelerated ageing and upregulation of Foxo or the process of autophagy for decelerated ageing. We supplement and strengthen this data by chemical-based intervention approach and by genetically inducing proteotoxic stress via expression of mutant Huntingtin protein that forms aggregates both locally in the LG and systemically in the fat body. Our results with multiple approaches demonstrate that LG homeostasis in terms of niche cell numbers, DNA damage accumulation and the extent of blood cell differentiation is abrogated in an accelerated ageing scenario. Our observations indicate that in the accelerated ageing model, the blood progenitors where we induce inflammaging display the typical features of ageing like loss of protein turnover, deregulation of autophagy and increase in ROS levels. We find that the cells undergoing accelerated ageing are capable of sensing this cellular stress and respond to it by switching on the Integrated Stress Response pathway. Genetic perturbation of ISR pathway components in the LG results in defective hematopoiesis. We hypothesized if sustained activation of the ISR pathway over and above existing levels of activation can restore blood cell homeostasis since mere switching on of the ISR pathway is not sufficient to restore blood cell homeostasis to a normal state in the accelerated ageing scenario in the LG. Our results indicate that genetic and chemical based approaches of activation of the ISR pathway in an accelerated ageing scenario are capable of restoring hematopoiesis. Taken together, our results provide novel mechanistic insights into the signalling biology underlying ageing at the cellular level during hematopoiesis.

Since the stem cell niche plays a crucial role in regulating homeostasis, we first investigated if the niche size is affected upon localized and systemic modulation of ageing. Our results show that the niche size reduces in the accelerated ageing scenario particularly upon Toll pathway activation in the niche, MZ progenitors or in the fat body. Now, it was earlier reported that Toll pathway hyperactivation in the niche leads to permeability barrier breakdown in the niche triggering hemocyte differentiation (Khadilkar et al., 2017). In the decelerated ageing conditions, Atg8 over-expression has no effect on the niche size whereas Foxo over-expression cell autonomously in the PSC, whole LG or systemically in the fat body reduces the niche size possibly because Foxo is a downstream effector and feedback regulator of insulin signaling (Piug et al., 2005) that attenuates insulin signaling (Ni et al., 2007). Insulin signalling plays an important role in regulating PSC size in the LG (Benmimoun et al., 2012) and Foxo over-expression dysregulates Insulin pathway (Ni et al., 2007). In the case of induction of accelerated ageing in the niche, there is increased plasmatocyte differentiation in both Toll and Imd activated background whereas crystal cell differentiation decreases upon Toll activation in the niche. Previous reports suggest that crystal cell lineage differentiation needs to be inhibited in order to promote lamellocyte differentiation making them mutually exclusive (Small et al., 2014, Diechsel et al., 2024). We observe a similar scenario here where Toll activation inhibits crystal cells promoting lamellocyte differentiation. However, it needs to be now tested if Toll pathway activation in the niche inhibits Notch activity promoting lamellocyte differentiation. Now in the decelerated ageing scenario, Atg8 over-expression has no effect on blood cell differentiation whereas Foxo inhibits crystal cell lineage, promotes plasmatocyte differentiation and shows lamellocyte differentiation in some LGs likely due to inhibition of Insulin signalling as is the case for example, for Insulin receptor (InR) knockdown in the niche that promotes plasmatocyte differentiation (Benmimoun et al., 2012). Domeless positive distal progenitor specific induction of accelerated ageing triggers the production of all lineages of hemocytes whereas induction of decelerated ageing suppresses crystal cell lineage in case of Foxo over-expression and promotes production of lamellocytes in few LGs whereas no effect on differentiation at all upon Atg8 over-expression. Intermediate zone-specific modulation of ageing via *chizGal4* behaves rather ideally wherein accelerated ageing triggers plasmatocyte, crystal cells and lamellocyte differentiation in the Imd activated background and a decrease in crystal cell differentiation in the Toll activated background possibly due to inhibition of Notch to promote lamellocyte production as has been discussed earlier. In the decelerated ageing background using *chizGal4*, there is no effect on hemocyte differentiation of all three lineages. The trend of increased plasmatocyte differentiation, decreased crystal cells and a modest induction of LGs with lamellocytes is replicated when we over-express Foxo in the differentiated blood cells using *hmlΔGal4* very similar to the phenotypes observed upon its over-expression in the niche or distal progenitors. Differentiated blood cell-specific induction of accelerated ageing by Imd activation results in increased production of all three differentiated lineages whereas Toll activation leads to complete suppression of plasmatocytes not only when activated in differentiated hemocytes but also in the intermediate progenitors that are transitioning towards terminal mature hemocytes. This could either mean that plasmatocytes made in these Toll activated conditions are a different subset of plasmatocytes that do not express the NimC1 antigen (encoded by NimRodC1 gene) since we now know that blood cells in *Drosophila* are very heterogeneous in nature (Fu et al., 2020, Cattenoz et al., 2020, Tattikota et al., 2020, Cho et al., 2020, Girard et al., 2021, Kharrat et al., 2022). In order to ascertain if this is a true phenotype we also checked if any of the parental transgenes are in *w^1118^*genetic background that was reported to have NimRodC1 mutation (Honti et al., 2013) but this was not the case indicating that these are true phenotypes. Differentiated hemocyte-specific activation of Toll again led to suppression of crystal cell lineage thereby promoting lamellocyte differentiation.

Systemic modulation of ageing is capable of influencing lymph gland hematopoiesis which is in line with previous literature demonstrating systemic regulation of hematopoiesis (Benmimoun et al., 2012, Shim et al., 2012, Yang et al., 2015, Cho et al., 2018, Koranteng et al., 2022). Now, loss of Atg8a in the muscles affects autophagy, reduces lifespan and muscle integrity and even leads to neuronal degeneration at organismal level (Xu et al., 2024). Similarly, Foxo over-expression in muscles (Demontis and Perrimon, 2010) and fat body (Hwangbo et al., 2004) was shown to extend lifespan at organismal level and remotely regulate aging in other tissues. Additionally, chronic Imd activation in the fat body resulted in shorter lifespan at an organismal level in the absence of infection (Sciambra & Chtarbanova., 2021). Based on these reports, we selected muscle and fat body as distant organs to investigate the inter-organ communication with the lymph gland and study the systemic effect of genetic modulation of cellular aging on the LG. Fat body-specific induction of accelerated ageing leads to an induction of all three hemocyte lineages in the LG whereas Atg8 over-expression has no significant effect on differentiation and Foxo over-expression in the fat body results in increased plasmatocyte differentiation whereas there is no effect on crystal cell or lamellocyte differentiation. Our results with muscle-specific *mhcGal4* show that accelerated ageing triggers differentiation whereas decelerated ageing suppresses differentiation or has no significant effect supporting our overall findings and indicating that both localized and systemic modulation of ageing is capable of altering hematopoiesis. DNA damage accumulation is one of the hallmarks of an aged cell. This phenomenon has been shown in an aged hematopoietic stem cell compartment as well (Beerman, 2017). We probed for γ-H2Ax that marks double strand breaks (DSBs) and our observations show that induction of accelerated ageing locally or systemically leads to DNA damage accumulation whereas decelerated ageing does not lead to such accumulation in line with previous literature revalidating our genetic models of cellular ageing.

We used alternative approaches to validate our findings using the expression of mutant Huntingtin locally in the niche and MZ progenitors in the LG or systemically in the fat body. In both cases there is reduction of niche size and an overall induction in differentiation of all three hemocyte lineages along with a decrease in the progenitor population when expressed in MZ which resembles the hallmark of stem cell exhaustion. We chose Huntington model as a medium to genetically induce proteotoxic stress based on previous studies that indicated that mutated huntingtin (HTT) protein in the Huntington’s autosomal dominant neurodegenerative disease is over-expressed and misfolds and is prone to aggregation and accumulation, thereby causing impairment in proteostasis (Koyuncu, 2017). In addition to this, we used a chemical intervention-based approach to accelerate or decelerate ageing. Bortezomib is an inhibitor of the ubiquitin-mediated proteasomal pathway (Chen et al., 2011) thereby resulting in increased protein instability, redox imbalance and accelerated aging (Manola et al., 2019). For decelerated ageing, we used Rapamycin as previous literature has shown that Rapamycin inhibits mTORC1 (mammalian Target of Rapamycin Complex1) and this inhibition of mTORC1 kinase by Rapamycin leads to upregulation of autophagy, increased resistance to starvation and lifespan extension in *Drosophila* (Bjedov et al., 2010). Our results show that Bortezomib induces whereas Rapamycin suppresses hemocyte differentiation very similar to our observations with the genetic modalities of cellular ageing. We used Rapamycin to investigate if it is capable of rescuing the Toll or Imd over-activation phenotypes in the LG. We find that Rapamycin is capable of rescuing the Imd activation phenotypes of increased plasmatocyte and crystal cell differentiation but not the lamellocyte differentiation phenotype. Rapamycin was unable to rescue the Toll over-activation phenotypes of increased differentiation which needs further investigation.

Since cellular stressors like impaired autophagy, elevated ROS levels, loss of protein turnover are sensed by signalling pathways like the Integrated Stress Response pathway (Kroemer et al., 2011, Costa-Matioli and Walter, 2020, Giardin et al., 2020, Ulfig and Jakob, 2024) we were curious to check if the ISR pathway is activated in the accelerated ageing background. We investigated *gcn2* transcript levels in blood cells upon modulation of ageing and also looked at peIF2α levels in the LG since it is an activation mark for the ISR pathway. Our analysis indicates that *gcn2* mRNA levels are higher in the accelerated ageing background consequently resulting in higher levels of peIF2α in the LG. We debated why blood cell homeostasis is not restored even upon ISR pathway activation in accelerated ageing background. To understand the intricacies of the role of ISR pathway components in the LG, we perturbed ISR pathway components in the hematopoietic progenitors and find that depletion of ISR pathway components results in increased blood cell differentiation whereas over-expression of *gcn2* or *atf4* has either no effect or suppresses hemocyte differentiation. We used whole animal mutants of ISR pathway components i.e. Gcn2 and Atf4 (*crc*). *gcn2* null or *crc* heterozygous mutants display increased differentiation of all three hemocyte lineages aligning with data obtained upon depletion of ISR components in hematopoietic progenitors. We chemically inhibited the ISR pathway using ISRIB, a known inhibitor of the ISR pathway (Zyryanova et al., 2021) and activated it using Histidinol (Taniuchi et al., 2016). ISRIB treatment induces hemocyte differentiation indicating that the ISR pathway is important for homeostasis. We then tested if accelerated ageing phenotypes upon Toll or Imd over-activation in progenitors are rescued by Histidinol treatment. Histidinol can more potently rescue Imd over-activation phenotypes whereas only plasmatocyte differentiation phenotype upon Toll over-activation can be rescued by Histidinol but not crystal cell or lamellocyte differentiation. The mechanistic link between Histidinol mediated ISR activation and Toll or Imd pathway warrant further investigation to understand how these NF-κB signalling pathways have an interplay with ISR. Lastly, since we debated that the existing activation levels of ISR cannot restore physiological hematopoiesis in an accelerated ageing scenario, we hypothesized if sustained activation of ISR over and above existing levels in accelerated ageing background can restore blood cell homeostasis. Genetic epistasis analysis shows that *gcn2* or *atf4* over-expression in *pirk* depletion (Imd activated) background in hematopoietic progenitors can restore the increased plasmatocyte or crystal cell differentiation whereas it cannot rescue the increased lamellocyte production. Furthermore, we also studied autophagy in these genetic backgrounds and found that the levels of autophagy are restored upon ectopic over-activation of ISR in Imd activated background.

Our results provide important mechanistic insights into the signalling biology underlying ageing wherein we show that ISR plays a critical role in combating stress especially in an inflammaging background and is crucial for ameliorating the detrimental effects of accelerated ageing. Our work paves way to investigate if aged hematopoietic stem cells and their niche micro-environment responds to age associated stressors similarly and whether the mechanisms bear similarity or not in the mammalian system. These findings will help in better understanding and calibrating risks associated with ARCH that have the potential to develop into hematological malignancies that are difficult to tackle increasingly due to challenges like chemoresistance. Taken together, organismal ageing associated changes impact at the cellular level that then has a spiralling down effect on the overall health and lifespan of an organism. Understanding the mechanisms and intricacies of cellular ageing will help in devising novel strategies to deal with disorders associated with ageing.

## Materials & Methods

### Drosophila Genetics

All the *Drosophila* stocks and crosses were maintained at 25°C, in a standard cornmeal diet containing corn starch & sugar as carbon source, malt extract containing trace number of vitamins and minerals, yeast extract as protein source and agar as solidifying agent. *Canton-S* was used as *wildtype* control. Tissue specific *Gal4* promoter line was used to drive the expression of *UAS* responder genes. Respective *UAS* or *Gal4* parent stocks or *Canton-S* were used as controls wherever appropriate. *Gal4* driver lines used were *collierGal4* driving *UAS- mCD8GFP* (gift from Dr. Michele Crozatier and Dr. Lucas Waltzer, Toulouse, France)*, tepIVGal4GFP* on Chr. II*, domeGal4GFP* on X*, hmlΔGal4GFP* on Chr. II (gift from Dr. Lucas Waltzer, Toulouse, France), *chizGal4GFP* on Chr. II (gifted by Dr. Bama Charan Mondal, BHU, India)*, dome-mesoGal4GFP* on Chr. III*, hmlGal4GFP* on Chr. II*, e33cGal4* on Chr. III (gifted by Dr. Maneesha Inamdar, JNCASR & DBT-InStem), *fbGal4* on Chr.III (Grönke et al., 2003)*, pplGal4* on Chr. II, (RRID:BDSC_58768) and muscle specific *mhcGal4* on Chr. III, (RRID:BDSC_55133) driving *UAS-mCD8GFP (*Chr. II, RRID: BL_5137). The *UAS*-transgene lines used were *UAS-pirkRNAi* (Chr. II, RRID: BL_67011)*, UAS-toll10B* (Chr. X, RRID: BDSC_58987)*, UAS-atg8GFP* (Chr. III, RRID:BDSC_51656)*, UAS-foxoGFP* (Chr. III, RRID: BDSC_43633), *UAS-HTT138Q.mRFP* (Chr. II, Gifted by Dr. Mayanglambam Dhruba Singh, NBRC)*, UAS-gcn2^act^, UAS-crcRA/cyO, crc^1/+^* on II*, gcn2^FRT^ ^12kb-/-^* on Chr III*, gcn2^wt^* rescue*; gcn2^FRT^ ^12kb-/-^* (Gifted by Dr. Hyung Don Ryoo, New York University Grossman School of Medicine, Kang et al., 2016)*, UAS-gcn2RNAi* (Chr. II, RRID: VDRC_103976), *UAS- eif2αRNAi* (Chr. II, RRID: VDRC_V104562), *UAS-atf4RNAi* (Chr. II, RRID: VDRC_109014).

### Antibodies

Antibodies used were mouse-raised anti-P1 (1:100, kind gift from Dr. Istvan Ando), mouse-raised anti-Hindsight (1:25, 1G9 – DSHB; RRID:Ab_528278), mouse-raised anti-Antp (1:25, 8C11- DSHB; RRID:Ab_528083), mouse-raised anti-γ2AX (1:200, UNC93-5.2.1 - DSHB; RRID:Ab_2618077), mouse-raised anti-Myospheroid (1:25, 6G11 - DSHB; RRID: Ab_528310), rabbit-raised anti-p-eIF2α (1:100, Cell Signaling technologies, RRID:Ab_119A11), CellROX Deep Red Reagent (2.5mM in DMSO, Invitrogen, C10422), PROTEOSTAT Protein detection assay (1:100, Enzo life sciences, CAT No.: ENZ-51023- KP050), rabbit-raised anti-P62/SQSTM1 (1:250, Proteintech, RRID:AB_10694431) and rabbit-raised anti-ATG8 (1:200, Sigma Aldrich, RRID:Ab_2939040) for immunofluorescence based experiments. Normal Goat Serum (HIMEDIA, RM10701) was used as the blocking agent. Alexa-Fluor 568 conjugated secondary antibodies – Goat-raised anti-Mouse 568 (1:400, Invitrogen, RRID: Ab_144696), Goat-raised anti-Mouse 633 (1:400, Invitrogen, RRID: AB_2535719) and Goat-raised anti-rabbit 568 (1:400, Invitrogen, RRID: AB_10563566) were used for immunofluorescence-based experiments.

### Lymph Gland Dissection, Immunohistochemistry & Mounting

Wandering late third instar larvae were used for lymph gland dissections. The dissections were performed in phosphate buffer saline (PBS), fixed in 4% paraformaldehyde for 20 minutes, followed by three washes with PBS containing 0.3% Triton-X (PBST) for 5 minutes each. The samples were then blocked in 20% normal goat serum for 20 minutes at room temperature followed by overnight primary antibody incubation at 4°C. This was followed by PBST washes, blocking, and treatment with appropriate Alexa-Fluor conjugated secondary antibody incubation for two hours at room temperature. This was again followed by three PBST washes for 5 minutes each. The LGs were then mounted in Vectashield mounting medium containing DAPI (Vector Laboratories, RRID: AB_2336790).

### Chemical treatment

For drug treatments, early to mid-third instar larvae were collected and transferred to vials without food containing few drops of distilled water (to prevent cuticle desiccation) and starved for 2 hours. The larvae were then transferred to food containing corresponding drugs to be used for treatment. Chemicals used include Rapamycin (40µg/ml, R0395, Sigma-Aldrich) dissolved in absolute ethanol, Bortezomib (10µM, 5.04314.2201, EMD Millipore) dissolved in DMSO (50ml, D8418, Sigma-Aldrich), ISRIB (5nm, SML0843, Sigma Aldrich) dissolved in DMSO & Histidinol (4mM, H6647, Sigma-Aldrich) dissolved in Nuclease free water (500 ml, 1097715, Invitrogen). For control, larvae were fed on food containing the respective solvent (vehicle) alone, post starvation. For each of the drug treatment experiments, at least 15 larvae were used for analysis. Lymph glands from the treated larvae for both the chemical and vehicle treatment were dissected 14 to 16 hours post treatment.

### Image acquisition and analysis of various Lymph Gland parameters

Images were acquired on Zeiss LSM 780 or Leica SP8 confocal microscope and analyzed for various hematopoietic parameters or for analysis of ROS levels, levels of autophagy, protein turn-over and p-eIF2ɑ levels. The details about the image analysis are included in the Supplemental Information.

### Quantitative Real-Time PCR

Details for the qRT-PCR experiment are included in the Supplemental Information

### Statistical Analysis

Immunofluorescence based experiments and their analysis was performed on at least 10 lymph glands dissected from wandering late third-instar larvae. The sample size for each of the experiments and the ‘N’ & ‘n’ values for all genotypes that denote the number of larvae & number of individual primary lymph gland lobes analysed per genotype respectively have been indicated in respective figure legends. Statistical analysis was performed using the GraphPad Prism Version 10 software (RRID:SCR_002798). For analysis of statistical significance, each experimental sample was tested with its respective control in a given experimental setup for all the data in each of the figures in order to estimate the P value. P values were determined by using a two-tailed unpaired Student’s t-test with Welch’s correction. Values are mean ± SD, and asterisks denote statistically significant differences (ns denotes not significant, * p < 0.05, ** p < 0.01, *** p < 0.001, **** p < 0.0001). Mutant genotypes were compared to the wildtype controls and the knockdown or over-expression genotypes were compared to their respective parental *Gal4* controls that were crossed to *wild-type* for all the statistical analysis performed. The experiments where chemical treatment was given were compared to the respective vehicle controls. No statistical method was used to pre-determine the sample size and the experiments were not randomized.

## Acknowledgements

We thank the Bloomington *Drosophila* Stock Center, Developmental Studies Hybridoma Bank and the fly community for fly stocks and antibodies. We would like to particularly thank Lucas Waltzer, Hyung Don Ryoo for various fly lines and reagents. We are thankful to the Stem Cell and Tissue Homeostasis lab for useful inputs and discussions.

## Author contributions

KG, RKI and RJK designed the experiments; KG, RKI, SS, MSI and JGL conducted the experiments; KG, RKI, SS analyzed the data; RJK procured the funding and resources and RKI, KG and RJK wrote the paper.

