## Supplemental Information for "Modulation of cellular ageing regulates Integrated Stress Response signalling to control blood cell homeostasis"

**Supplemental Methods:**

**Genotypes and genetic crosses:**

**Figure 1:** To evaluate the conventional hallmarks of ageing including estimation of autophagic flux, ROS levels & proteostasis regulation via protein turnover assessment in the *tepIV* - positive core progenitors, the following crosses were set up - For autophagic flux assessment, *tepIVGal4GFP x UAS-pirkRNAi* (accelerated ageing) & *tepIVGal4GFP x UAS-atg8GFP* (decelerated ageing) was compared with *tepIVGal4GFP x wt as wildtype* control. For ROS levels estimation, *tepIVGal4GFP x UAS-toll10B* (accelerated ageing), *tepIVGal4GFP x UAS-atg8GFP* & *tepIVGal4GFP x UAS-foxoGFP* (decelerated ageing) were compared with *tepIVGal4GFP x wt as wildtype* control. For assessment of proteostasis via protein turnover estimation, *tepIVGal4GFP x UAS-pirkRNAi* & *tepIVGal4GFP x UAS-toll10B* (accelerated ageing), *tepIVGal4GFP x UAS-atg8GFP* & *tepIVGal4GFP x UAS-foxoGFP* (decelerated ageing) were compared to *tepIVGal4GFP x wt* as *wildtype* control.

**Figure 2:** The following crosses were set up to study the effect of modulation of cellular ageing on PSC niche size by localised or systemic cellular subsets of lymph gland (LG):

a. **PSC Niche - specific:** *collierGal4GFP x UAS-pirkRNAi* & *collierGal4GFP x UAS-toll10B* (accelerated ageing), *collierGal4GFP x UAS-atg8GFP* & *collierGal4GFP x UAS-foxoGFP* (decelerated ageing) and *collierGal4GFP x wt* (Control).
b. **Distal progenitor – specific:** *domeGal4GFP x UAS-pirkRNAi* & *domeGal4GFP x UAS-toll10B* (accelerated ageing), *domeGal4GFP x UAS-atg8GFP* & dome*Gal4GFP x UAS-foxoGFP* (decelerated ageing) and *domeGal4GFP x wt* (Control).
c. **Cortical zone differentiated hemocyte – specific:** *hmlΔGal4GFP* x *UAS-pirkRNAi* & *hmlΔGal4GFP x UAS-toll10B* (accelerated ageing), *hmlΔGal4GFP x UAS-atg8GFP* & *hmlΔGal4GFP x UAS-foxoGFP* (decelerated ageing) and *hmlΔGal4GFP x wt* (Control).
d. **Whole LG – specific:** *e33cGal4 x UAS-pirkRNAi* & *e33cGal4 x UAS-toll10B* (accelerated ageing), *e33cGal4 x UAS-atg8GFP* & *e33cGal4 x UAS-foxoGFP* (decelerated ageing) and *e33cGal4 x wt* (Control).
e. **Systemic fat body – specific:** *fbGal4 x UAS-pirkRNAi* & *fbGal4 x UAS-toll10B* (accelerated ageing), *fbGal4 x UAS-atg8GFP* & *fbGal4 x UAS-foxoGFP* (decelerated ageing) and *fbGal4 x wt* as *wildtype* control.

**Figure 3:** The following crosses were set up to study the effect of genetic modulation of cellular ageing in distal-progenitor population using *domeGal4* on lymph gland hematopoiesis by inducing accelerated ageing (*domeGal4GFP* x *UAS-pirkRNAi and domeGal4GFP* x *UAS-toll10B*) and decelerated ageing (*domeGal4GFP* x *UAS-atg8GFP and domeGal4GFP* x *UAS-foxoGFP*) as compared to *domeGal4GFP x wt* as *wildtype* control.

**Figure 4:** The following crosses were set up to study the effect of genetic modulation of cellular ageing systemically in muscles using *mhc-Gal4* on lymph gland hematopoiesis by inducing accelerated ageing (*mhcGal4GFP* x *UAS-pirkRNAi and mhcGal4GFP* x *UAS-toll10B*) and decelerated ageing (*mhcGal4GFP* x *UAS-atg8GFP and mhcGal4GFP* x *UAS-foxoGFP*) as compared to *mhcGal4GFP* x *wt* as *wildtype* control.

**Figure 5:** The following crosses were set up to study the effect of inducing proteotoxic stress via *HTT138Q.mRFP* expression in PSC (using *collierGal4*) or core progenitors (using *tepIVGal4*). For niche, *collierGal4GFP x UAS-HTT138Q.mRFP was compared with collierGal4GFP x wt as wildtype control. For progenitors, tepIVGal4GFP x UAS-HTT138Q.mRFP was compared with tepIVGal4GFP x wt as wildtype control.*

**Figure 6:** The following crosses were set up to study the systemic effect of chemical modulation of ageing on lymph gland hematopoiesis upon treatment with Bortezomib (causing accelerated ageing) on *tepIVGal4GFP x wt* larvae and compared with its solvent (vehicle) control. Similarly, Rapamycin treatment (causing decelerated aging) on *wildtype* larvae (*tepIVGal4GFP x wt)* and in an accelerated ageing scenario where Toll or Imd is activated in core progenitors to investigate its effect on rescue of differentiation (*tepIVGal4GFP x UAS-pirkRNAi and tepIVGal4GFP x UAS-toll10B)* when compared with its solvent (vehicle) control.

**Figure 7:** The following crosses were set up to check for the expression levels of *gcn2* in the hemocyte population (using *hmlGal4*) & *p-eIF2α* in core-progenitors (using *tepIVGal4*): For estimation of *gcn2* mRNA transcript levels in all hemocytes, following crosses were set up: *hmlGal4GFP* x *UAS-pirkRNAi & hmlGal4GFP x UAS-toll10B* (accelerated ageing), *hmlGal4GFP x UAS-atg8GFP & hmlGal4GFP x UAS-foxoGFP* (decelerated ageing) were compared to *hmlGal4GFP x wt* as *wildtype* control and for estimation of *p-eIF2α* in core-progenitors, *tepIVGal4GFP x UAS-pirkRNAi & tepIVGal4GFP x UAS-toll10B* (accelerated ageing), *tepIVGal4GFP x UAS-atg8GFP* & *tepIVGal4GFP x UAS-foxoGFP* (decelerated ageing) were compared to *tepIVGal4GFP x wt* as *wildtype* control. Genetic modulation of ISR pathway via knockdown (*UAS-gcn2RNAi, UAS-eIF2αRNAi, UAS-atf4RNAi*) or overexpression (*UAS-gcn2$^{act}$* & *UAS-crcRA)* of ISR components in *tepIVGal4* - positive core progenitors and its effect on blood cell differentiation in lymph gland was studied by setting up the following crosses: *tepIVGal4GFP x UAS-gcn2RNAi, tepIVGal4GFP x UAS-eIF2αRNAi, tepIVGal4GFP x UAS-atf4RNAi, tepIVGal4GFP x UAS-gcn2$^{act}$, tepIVGal4GFP x UAS-crcRA* as compared to *tepIVGal4GFP x wt* as *wildtype* control.

**Figure 8:** ISR components including *gcn2$^{act}$* & *crcRA* were ectopically over-expressed in distal-progenitor population using *dome-mesoGal4GFP* in IMD pathway upregulated (*pirkRNAi*) background (Resultant genotype: *UAS-gcn2$^{act}$ or UAS-crcRA/UAS-pirkRNAi; dome-mesoGal4GFP/dome-mesoGal4GFP or dome-mesoGal4GFP/TM6B*) and compared with *dome-mesoGal4GFP x UAS-pirkRNAi* as control to investigate blood cell homeostasis and status of autophagy upon ISR activation in accelerated ageing genetic background.

**Figure S1:** The following crosses were set up to study the effect of modulation of cellular ageing on blood cell differentiation upon induction of accelerated ageing (*UAS-pirkRNAi & UAS-toll10B*) and decelerated ageing (*UAS-atg8GFP & UAS-foxoGFP*) in PSC niche using *collierGal4 - collierGal4GFP x UAS-pirkRNAi*, *collierGal4GFP x UAS-toll10B*, *collierGal4GFP x UAS-atg8GFP*, *collierGal4GFP x UAS-foxoGFP* and *collierGal4GFP x wt* as *wildtype* control.

**Figure S2:** The following crosses were set up to study the effect of modulation of cellular ageing on hemocyte differentiation in lymph gland upon induction of accelerated ageing (*UAS-pirkRNAi & UAS-toll10B*) and decelerated ageing (*UAS-atg8GFP & UAS-foxoGFP*) in intermediate-progenitor population using *chizGal4*: *chizGal4GFP x UAS-pirkRNAi*, *chizGal4GFP x UAS-toll10B*, *chizGal4GFP x UAS-atg8GFP*, *chizGal4GFP x UAS-foxoGFP* and *chizGal4GFP x wt* as *wildtype* control.

**Figure S3:** The following crosses were set up to study the effect of modulation of cellular ageing on lymph gland hematopoiesis upon induction of accelerated ageing (*UAS-pirkRNAi & UAS-toll10B*) and decelerated ageing (*UAS-atg8GFP & UAS-foxoGFP*) in cortical zone specific - differentiated hemocyte population using *hmlΔGal4: hmlΔGal4GFP* x *UAS-pirkRNAi*, *hmlΔGal4GFP x UAS-toll10B*, *hmlΔGal4GFP x UAS-atg8GFP*, *hmlΔGal4GFP x UAS-foxoGFP* and *hmlΔGal4GFP x wt* as *wildtype* control.

**Figure S4:** The following crosses were set up to study the effect of systemic modulation of cellular ageing on lymph gland hematopoiesis upon induction of accelerated ageing (*UAS-pirkRNAi & UAS-toll10B*) and decelerated ageing (*UAS-atg8GFP & UAS-foxoGFP*) in fat body using *pplGal4: pplGal4 x UAS-pirkRNAi*, *pplGal4 x UAS-toll10B*, *pplGal4 x UAS-atg8GFP*, *pplGal4 x UAS-foxoGFP* and *pplGal4 x wt* as *wildtype* control.

**Figure S5:** The following crosses were set up to study the effect of modulation of cellular ageing on DNA damage in localised or systemic cellular subsets of lymph gland (LG):

a. PSC niche - specific *collierGal4GFP x UAS-pirkRNAi* (accelerated ageing), *collierGal4GFP* x *UAS-foxoGFP* (decelerated aging) and *collierGal4GFP* x *wt* (Control).
b. Distal progenitor – specific *domeGal4GFP* x *UAS-pirkRNAi* (accelerated ageing), dome*Gal4GFP* x *UAS-foxoGFP* (decelerated ageing) and *domeGal4GFP* x *wt* (Control).
c. Cortical zone differentiated hemocyte – specific *hmlΔGal4GFP x UAS-pirkRNAi* (accelerated ageing)*, hmlΔGal4GFP x UAS-foxoGFP* (decelerated ageing) *and hmlΔGal4GFP x wt* (Control).
d. Whole LG – specific *e33cGal4* x *UAS-pirkRNAi* (accelerated ageing)*, e33cGal4 x UAS-foxoGFP* (decelerated ageing) and *e33cGal4* x *wt* (Control).
e. Systemic fat body – specific *fbGal4* x *UAS-pirkRNAi* (accelerated ageing), *fbGal4* x *UAS-foxoGFP* (decelerated ageing) and *fbGal4* x *wt* as *wildtype* control.

**Figure S6:** The following crosses were set up to study the systemic effect of proteotoxic stress via *HTT138Q.mRFP* expression in the fat body on LG homeostasis: *fbGal4* x *UAS-HTT138Q.mRFP* was compared with *fbGal4* x *wt* as *wildtype* control.

**Figure S7:** To study the effect of ISR mutants on hemocyte differentiation in lymph gland, the following lines were used: *gcn2* $^{FRT\ 12kb\ -/-}$ null mutant & *gcn2*$^{wt}$ rescue*; gcn2*$^{FRT\ 12kb\ -/-}$ as its control and *crc*$^{1/+}$ hypomorphic allele & a *wildtype* control.

**Figure S8:** The following crosses were set up to study the effect of chemical modulation of ISR pathway on lymph gland homeostasis upon treatment with ISRIB on *tepIVGal4GFP x wt* larvae compared with its solvent (vehicle) control. Similarly, Histidinol treatment under accelerated ageing background (*tepIVGal4GFP x UAS-pirkRNAi & tepIVGal4GFP x UAS-*

*toll10B)* was performed to investigate its effect on rescue of differentiation and compared with its solvent (vehicle) control.

**Image analysis:**

**a. Hematopoietic parameters:**

Confocal images were captured using either Zeiss LSM 780 or Leica SP8 confocal microscope. Z projection of the confocal images was used for estimating various lymph gland parameters using ImageJ/Fiji software. Plasmatocyte differentiation Index was estimated by measuring the percentage of P1 positive area divided by the total area of the lymph gland primary lobe. The prohemocyte Index was estimated by measuring the percentage of *tepIV-GFP* or *dome-GFP* positive area divided by the total area of lymph gland primary lobe. Freehand selection tool was used for measuring the area of the plasmatocytes or the prohemocytes. For the quantitation of Antp, Hnt, and γH2AX, the positive signals for respective markers were manually counted using the multipoint tool. For quantitation of Mys, the lymph glands were categorised as positive or negative based on presence or absence of lamellocytes and percentage of lamellocyte positive lymph glands was estimated. The lymph gland quantitations were done for individual primary lymph gland lobes.

**b. Analysis of expression:**

Confocal images were captured using either Zeiss LSM 780 or Leica SP8 confocal microscope. For estimation of ROS levels using CellROX Deep Red Reagent in *tepIVGal4*-specific core-progenitor population in the lymph gland using ImageJ/Fiji software (RRID:SCR_003070), the images for *tepIVGal4GFP* x *wt, tepIVGal4GFP* x *UAS-toll10B* (accelerated aging)*, tepIVGal4GFP* x *UAS-atg8GFP & tepIVGal4GFP* x *UAS-foxoGFP* (decelerated aging) were acquired at the same intensity settings/parameters and mean fluorescent intensity (represented as arbitrary units) in the progenitors was calculated keeping the threshold value of fluorescent intensity uniform as that of *tepIVGal4GFP* x *wt* for all the above crosses. Similar protocol was followed for estimating the mean fluorescence intensity of p-eIF2α in *tepIVGal4GFP* x *wt* and *tepIVGal4GFP* x *UAS-pirkRNAi, tepIVGal4GFP* x *UAS-toll10B* (accelerated ageing)*, tepIVGal4GFP* x *UAS-atg8GFP and tepIVGal4GFP* x *UAS-foxoGFP* (decelerated ageing) which was calculated keeping the threshold value of fluorescent intensity uniform as that of *tepIVGal4GFP* x *wt* for all the above crosses.

For the quantitation of proteotoxic stress and autophagic flux, we unbiasedly selected several fields from each image and within each field we counted the total number of punctae (p) for Proteostat, p62 & Atg8 respectively and also counted the total number of nuclei (n). In our quantitation, we took the ratio of p/n which is representative of puncta per cell considering all lymph gland cells are uninucleate. Thus, we compared the Proteostat or p62 or Atg8 puncta per cell in the lymph gland of different experimental sets.

**Quantitative Real Time PCR:**

Hemolymph was extracted from 300 larvae mounted in cold PBS by puncturing the cuticle using minutien insect pins. The hemolymph was pelleted by centrifuging at 2000g for 7 minutes at 4°C. The supernatant was removed and the hemolymph pellet was lysed in TRIzol (Ambion

– life technologies, Catalogue No.: 11596018). The lysates were stored at -80°C. Hemolymph collection were done in batches of 50 to 100 larvae and once the hemolymph from all 300 larvae were done, RNA was isolated by pooling all the aqueous layers post-chloroform treatment, followed by RNA isolation according to the manufacturer's protocol. RNA yield was quantified using Nanodrop. 1µg of mRNA was reverse transcribed using oligo-dT primers (Promega, C110A) and ImProm-II (Promega, A3800). Quantitation of the mRNA transcripts was done using SYBR green chemistry (Thermo Fisher Scientific, Catalogue No.: 4367659) in the Quantstudio 5 RT PCR system (Thermo Fisher Scientific) in quadruplets of 10µl reaction. The data was analyzed using the ΔΔCt method and relative mRNA expression was normalized to *rp49*. Fold change calculations were done in comparison to *wildtype* control. The experiment was done in biological triplicates & statistical analysis was performed using One-way ANOVA (Dunnett) for comparison of all test genotypes with *wildtype* control genotype.

**List of Primers:**

qRT primers in 5' to 3' direction

| | | |
|---|---|---|
| *gcn2* | Forward | CCAACGGACATACGGATACAAC |
| *gcn2* | Reverse | CGTAGCTCTTGGGATTGAGCC |
| *rp49* | Forward | GCTAAGCTGTCGCACAAATG |
| *rp49* | Reverse | GTTCGATCCGTAACCGATGT |

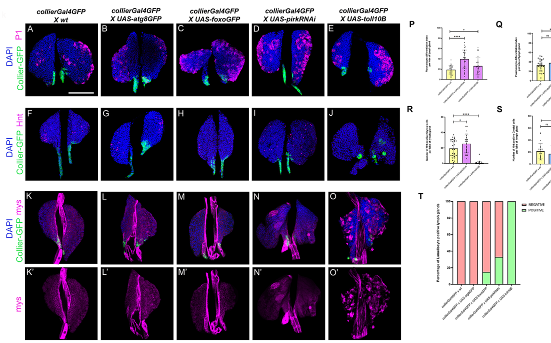

collierGal4GFP X wt
collierGal4GFP X UAS-atg8GFP
collierGal4GFP X UAS-foxoGFP
collierGal4GFP X UAS-pirkRNAi
collierGal4GFP X UAS-toll10B
A
B
C
D
E
DAPI
Collier-GFP
P1
F
G
H
I
J
Hnt
K
L
M
N
O
mys
K'
L'
M'
N'
O'
P
Q
R
S
T
Plasmatocyte differentiation index per lobe of lymph gland
Number of Hnt positive Crystal cells per lobe of lymph gland
collierGal4GFP x wt
collierGal4GFP x UAS-pirkRNAi
collierGal4GFP x UAS-toll10B
ns
Percentage of Lamellocyte positive lymph glands
NEGATIVE
POSITIVE
collierGal4GFP x wt
collierGal4GFP x UAS-atg8GFP
collierGal4GFP x UAS-foxoGFP
collierGal4GFP x UAS-pirkRNAi
collierGal4GFP x UAS-toll10B

**Supplementary Figure Legends**

**Supplementary Figure 1: PSC niche-specific modulation of cellular ageing affects lymph gland homeostasis**

Plasmatocyte differentiation marked by P1 (magenta) or crystal cell differentiation marked by Hindsight (Hnt) (magenta) or lamellocyte differentiation marked by β-integrin-specific Myospheroid (Mys) (magenta) upon PSC niche-specific (using *collierGal4*) expression of *UAS-atg8GFP* (Fig. S1B, G, L), *UAS-foxoGFP* (Fig. S1C, H, M), *UAS-pirkRNAi* (Fig. S1D, I, N) or *UAS-toll10B* (Fig. S1E, J, O) as compared to *wildtype* control (Fig. S1A, F, K). GFP expression (green) is driven by *collierGal4* (Fig. S1A-O). Nuclei are stained with DAPI (Blue). Graphical representation of plasmatocyte differentiation index or number of crystal cells or percentage of lamellocyte positive lymph glands upon *collierGal4* mediated expression of *UAS-pirkRNAi* or *UAS-toll10B* (Fig. S1P, R, T) or *UAS-atg8GFP* or *UAS-foxoGFP* (Fog. S1Q, S, T) as compared to *wildtype* control. For Plasmatocyte differentiation in accelerated ageing scenario: *collierGal4 x UAS-pirkRNAi* (N = 16 , n = 31) and *collierGal4 x UAS-toll10B* (N = 16, n = 31) as compared to *collierGal4 x wt* (N = 14, n = 27) and in decelerated ageing scenario: *collierGal4 x UAS-atg8GFP* (N = 12, n = 23), *collierGal4 x UAS-foxoGFP* (N = 15 , n = 30) as compared to *collierGal4 x wt* (N = 13 , n = 25). For crystal cell numbers in accelerated ageing scenario: *collierGal4 x UAS-pirkRNAi* (N = 16 , n = 31) and *collierGal4 x UAS-toll10B* (N = 20, n = 40) as compared to *collierGal4 x wt* (N = 14, n = 27) and in decelerated ageing scenario: *collierGal4 x UAS-atg8GFP* (N = 13, n = 26), *collierGal4 x UAS-foxoGFP* (N = 14 , n = 28) as compared to *collierGal4 x wt* (N = 11, n = 22). N denotes the number of larvae & n denotes the number of individual primary lymph gland lobes analysed per genotype. Individual data points in the graphs represent individual primary lobes of the Lymph gland. Values are mean ± SD, and asterisks denote statistically significant differences (ns denotes not significant, *$p<0.05$, **$p<0.01$, *** $p<0.001$, ****$p<0.0001$). Student's t-test with Welch's correction was performed for the statistical analysis. Scale Bar: 50 µm (A-O').

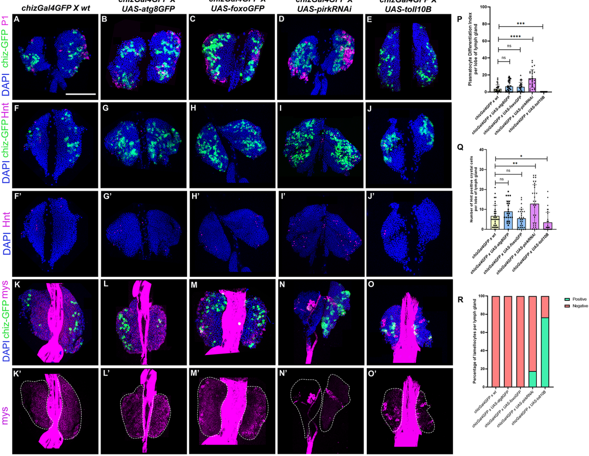

chizGal4GFP X wt
chizGal4GFP X UAS-atg8GFP
chizGal4GFP X UAS-foxoGFP
chizGal4GFP X UAS-pirkRNAi
chizGal4GFP X UAS-toll10B
DAPI chiz-GFP P1
DAPI chiz-GFP Hnt
DAPI Hnt
DAPI chiz-GFP mys
mys
A
B
C
D
E
F
G
H
I
J
F'
G'
H'
I'
J'
K
L
M
N
O
K'
L'
M'
N'
O'
P
Plasmatocyte Differentiation Index per lobe of lymph gland
ns
ns
****
***
Q
Number of Hnt positive crystal cells per lobe of lymph gland
ns
ns
**
*
R
Percentage of lamellocytes per lymph gland
Positive
Negative
chizGal4GFP x wt
chizGal4GFP x UAS-atg8GFP
chizGal4GFP x UAS-foxoGFP
chizGal4GFP x UAS-pirkRNAi
chizGal4GFP x UAS-toll10B

**Supplementary Figure 2: Intermediate progenitor-specific modulation of cellular ageing alters blood cell differentiation in the LG**

Plasmatocyte differentiation marked by P1 (magenta) or crystal cell differentiation marked by Hindsight (Hnt, magenta) or lamellocyte differentiation marked by β-integrin-specific Myospheroid (Mys, (magenta) upon intermediate progenitor-specific (using *chizGal4*) expression of *UAS-atg8GFP* (Fig. S2B, G, L), *UAS-foxoGFP* (Fig. S2C, H, M), *UAS-pirkRNAi* (Fig. S2D, I, N) or *UAS-toll10B* (Fig. S2E, J, O) as compared to *wildtype* control (Fig. S2A, F, K). GFP expression (green) is driven by *chizGal4* (Fig. S2A-O). Nuclei are stained with DAPI (Blue). Graphical representation of plasmatocyte differentiation index (Fig. S2P) or number of crystal cells (Fig. S2Q) or percentage of lamellocyte positive lymph glands (Fig. S2R) upon *chizGal4* mediated expression of *UAS-pirkRNAi*, *UAS-toll10B*, *UAS-atg8GFP* or *UAS-foxoGFP* as compared to *wildtype* control. For Plasmatocyte differentiation, *chizGal4 x UAS-pirkRNAi* (N = 14, n = 28), *chizGal4 x UAS-toll10B* (N = 15, n = 30), *chizGal4 x UAS-atg8GFP* (N = 14, n = 28), *chizGal4 x UAS-foxoGFP* (N = 15, n = 30) were analyzed as compared to *chizGal4 x wt* (N = 15, n = 30). For crystal cell numbers, *chizGal4 x UAS-pirkRNAi* (N = 16, n = 32), *chizGal4 x UAS-toll10B* (N = 16, n = 32), *chizGal4 x UAS- atg8GFP* (N = 14, n = 27), *chizGal4 x UAS-foxoGFP* (N = 15, n = 30) were analyzed as compared to *chizGal4 x wt* (N = 18, n = 35). For lamellocyte differentiation, *chizGal4 x UAS- pirkRNAi* (N = 16), *chizGal4 x UAS-toll10B* (N = 14), *chizGal4 x UAS-atg8GFP* (N = 19), *chizGal4 x UAS-foxoGFP* (N= 16) were analyzed as compared to *chizGal4 x wt* (N = 15). N denotes the number of larvae & n denotes the number of individual primary lymph gland lobes analyzed per genotype. Individual data points in the graphs represent individual primary lobes of the Lymph gland. Values are mean ± SD, and asterisks denote statistically significant differences (ns denotes not significant, $^*p<0.05$, $^{**}p<0.01$, $^{***}p<0.001$, $^{****}p<0.0001$). Student's t-test with Welch's correction was performed for the statistical analysis. Scale Bar: 50 µm (A-O').

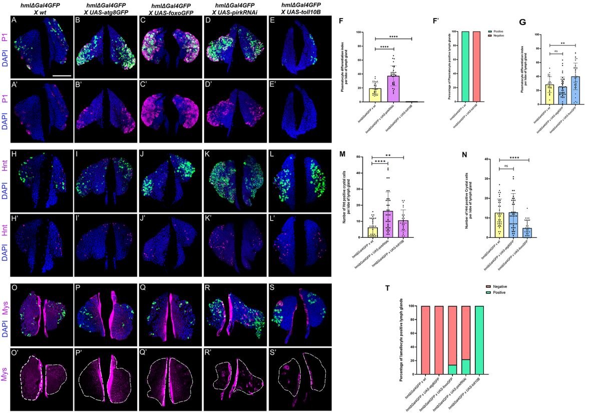
hmlΔGal4GFP X wt
hmlΔGal4GFP X UAS-atg8GFP
hmlΔGal4GFP X UAS-foxoGFP
hmlΔGal4GFP X UAS-pirkRNAi
hmlΔGal4GFP X UAS-toll10B
hmlΔ-GFP P1 DAPI
P1 DAPI
hmlΔ-GFP Hnt DAPI
Hnt DAPI
hmlΔ-GFP Mys DAPI
Mys
A B C D E
A' B' C' D' E'
H I J K L
H' I' J' K' L'
O P Q R S
O' P' Q' R' S'
F
Plasmatocyte differentiation index per lobe of lymph gland
hmlΔGal4GFP x wt
hmlΔGal4GFP x UAS-pirkRNAi
hmlΔGal4GFP x UAS-toll10B
F'
Percentage of Plasmatocyte positive lymph glands
Positive
Negative
G
Plasmatocyte differentiation index per lobe of lymph gland
hmlΔGal4GFP x UAS-atg8GFP
hmlΔGal4GFP x UAS-foxoGFP
ns
M
Number of Hnt positive crystal cells per lobe of lymph gland
N
Number of Hnt positive Crystal cells per lobe of lymph gland
T
Percentage of lamellocyte positive lymph glands

**Supplementary Figure 3: Cortical zone differentiated hemocyte – specific modulation of cellular ageing cell autonomously regulates blood cell differentiation**

Plasmatocyte differentiation marked by P1 (magenta) or crystal cell differentiation marked by Hindsight (Hnt in magenta) or lamellocyte differentiation marked by β-integrin-specific Myospheroid (Mys in magenta) upon cortical zone differentiated hemocyte (using *hmlΔGal4*) mediated expression of *UAS-atg8GFP* (Fig. S3B - B', I – I', P – P'), *UAS-foxoGFP* (Fig. S3C – C', S3J – J', S3Q – Q'), *UAS-pirkRNAi* (Fig. S3D – D', S3K – K', S3R – R') or *UAS-toll10B* (Fig. S3E – E', S3L – L', S3S – S') as compared to *wildtype* control (S3A – A', S3H – H', S3O – O'). GFP (green) is driven by *hmlΔGal4* which marks differentiated hemocytes in the cortical zone (Fig. S3A – E, S3H – L, S3O - S). Nuclei are stained with DAPI (Blue). Graphical representation of plasmatocyte differentiation index in accelerated ageing (Fig. S3F-F') and decelerated ageing scenario (Fig. S3G) or number of crystal cells in accelerated (Fig. S3M) and decelerated ageing scenario (Fig. S3N) or percentage of lamellocyte positive lymph glands (Fig. S3T) as compared to *wildtype* control. For plasmatocyte differentiation, decelerated aging: *hmlΔGal4 x UAS-atg8GFP* (N = 13, n = 26) & *hmlΔGal4 x UAS-foxoGFP (*N = 13, n = 26) as compared to *wildtype* control (N = 14, n = 27), accelerated aging: *hmlΔGal4 x UAS-pirkRNAi (*N = 16, n = 32), *hmlΔGal4 x UAS-toll10B (*N = 10, n = 20) as compared to *wildtype* control (N = 14, n = 28). For crystal cell differentiation, decelerated aging: *hmlΔGal4 x UAS-atg8GFP* (N = 13, n = 26) & *hmlΔGal4 x UAS-foxoGFP (*N = 11, n = 21) as compared to *wildtype* control (N = 17, n = 34), accelerated aging: *hmlΔGal4 x UAS-pirkRNAi (*N = 17, n = 33), *hmlΔGal4 x UAS-toll10B (*N = 12, n = 24) as compared to *wildtype* control (N = 15, n = 30). For lamellocyte differentiation, *hmlΔGal4 x UAS-atg8GFP* (N = 15), *hmlΔGal4 x UAS-foxoGFP (*N = 15), *hmlΔGal4 x UAS-pirkRNAi (*N = 15), *hmlΔGal4 x UAS-toll10B (*N = 15) as compared to *wildtype* control *hmlΔGal4 x wt* (N = 15). N denotes the number of larvae & n denotes the number of individual primary lymph gland lobes analysed per genotype. Individual data points in the graphs represent individual primary lobes of the Lymph gland. Values are mean ± SD, and asterisks denote statistically significant differences (ns denotes not significant, $^{**}p < 0.01$, $^{***}p < 0.001$, $^{****}p < 0.0001$). Student's t-test with Welch's correction was performed for the statistical analysis. Scale Bar: 50 µm (A-S').

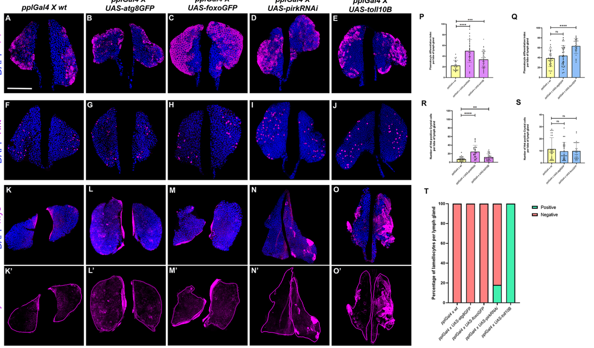
A
B
C
D
E
F
G
H
I
J
K
L
M
N
O
K'
L'
M'
N'
O'
P
Q
R
S
T
pplGal4 X wt
pplGal4 X UAS-atg8GFP
pplGal4 X UAS-foxoGFP
pplGal4 X UAS-pirkRNAi
pplGal4 X UAS-toll10B
P1
DAPI
Hnt
DAPI
mys
DAPI
mys
Plasmatocyte differentiation index per lobe of lymph gland
Number of Hnt positive Crystal cells per lobe of lymph gland
****
***
**
ns
Percentage of lamellocytes per lymph gland
Positive
Negative
pplGal4 x wt
pplGal4 x UAS-atg8GFP
pplGal4 x UAS-foxoGFP
pplGal4 x UAS-pirkRNAi
pplGal4 x UAS-toll10B

**Supplementary Figure 4: Fat body-specific systemic modulation of ageing affects blood cell homeostasis in the LG.**

Plasmatocyte differentiation marked by P1 (magenta) or crystal cell differentiation marked by Hindsight (Hnt, magenta) or lamellocyte differentiation marked by β-integrin-specific Myospheroid (Mys, magenta) upon systemic fat body-specific (using *pplGal4*) expression of *UAS-atg8GFP* (S4B, G, L), *UAS-foxoGFP* (S4C, H, M), *UAS-pirkRNAi* (S4D, I, N) or *UAS-toll10B* (S4E, J, O) as compared to *wildtype* control (S4A, F, K). Nuclei are stained with DAPI (Blue). Graphical representation of plasmatocyte differentiation index or number of crystal cells or percentage of lamellocyte positive lymph glands upon *pplGal4* mediated expression of *UAS-pirkRNAi* or *UAS-toll10B* (S4P, R, T) or *UAS-atg8GFP* or *UAS-foxoGFP* (S4Q, S, T) as compared to wildtype control. For Plasmatocyte differentiation in accelerated ageing scenario: *pplGal4* x *UAS-pirkRNAi* (N = 12, n = 24) and *pplGal4 x UAS-toll10B* (N = 21, n = 42) as compared to *pplGal4 x wt* (N = 11, n = 22) and in decelerated ageing scenario: *pplGal4 x UAS-atg8GFP* (N = 16, n = 31), *pplGal4 x UAS-foxoGFP* (N = 15, n = 30) as compared to *pplGal4 x wt* (N = 16, n = 32). For crystal cell numbers in accelerated ageing scenario: *pplGal4 x UAS-pirkRNAi* (N = 12, n = 23) and *pplGal4 x UAS-toll10B* (N = 16, n = 32) as compared to *pplGal4 x wt* (N = 13, n = 26) and in decelerated ageing scenario: *pplGal4 x UAS-atg8GFP* (N = 14, n = 27), *pplGal4 x UAS-foxoGFP* (N = 15, n = 29) as compared to *pplGal4 x wt* (N = 16, n = 31). A minimum of 15 larvae were analyzed for lamellocyte differentiation per genotype. N denotes the number of larvae & n denotes the number of individual primary lymph gland lobes analysed per genotype. Individual data points in the graphs represent individual primary lobes of the Lymph gland. Values are mean ± SD, and asterisks denote statistically significant differences (ns denotes not significant, **$p<0.01$, *** $p<0.001$, ****$p<0.0001$). Student's t-test with Welch's correction was performed for the statistical analysis. Scale Bar: 50 µm (A-O').

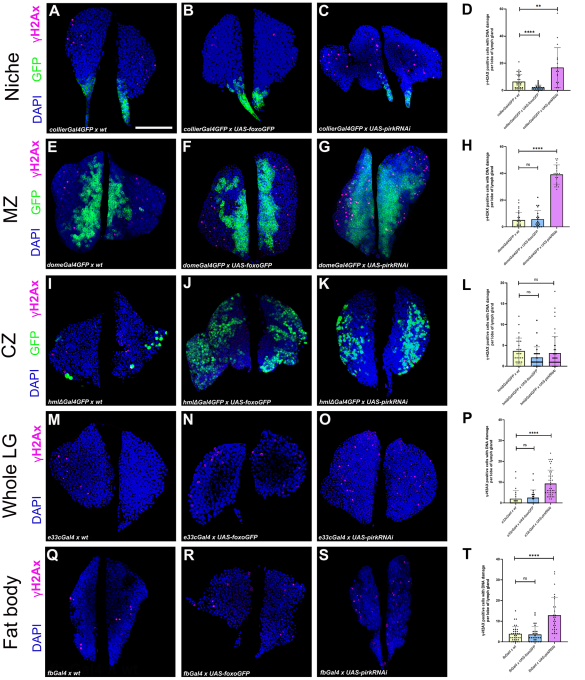
Niche
MZ
CZ
Whole LG
Fat body
γH2Ax
GFP
DAPI
A
B
C
D
E
F
G
H
I
J
K
L
M
N
O
P
Q
R
S
T
collierGal4GFP x wt
collierGal4GFP x UAS-foxoGFP
collierGal4GFP x UAS-pirkRNAi
domeGal4GFP x wt
domeGal4GFP x UAS-foxoGFP
domeGal4GFP x UAS-pirkRNAi
hmlΔGal4GFP x wt
hmlΔGal4GFP x UAS-foxoGFP
hmlΔGal4GFP x UAS-pirkRNAi
e33cGal4 x wt
e33cGal4 x UAS-foxoGFP
e33cGal4 x UAS-pirkRNAi
fbGal4 x wt
fbGal4 x UAS-foxoGFP
fbGal4 x UAS-pirkRNAi
γ-H2AX positive cells with DNA damage per lobe of lymph gland
**
****
ns

**Supplementary Figure 5: LG – specific or systemic modulation of cellular ageing regulates DNA damage accumulation in the LG.**

Cellular subsets of various lymph gland subsets undergoing DNA damage marked by *ɣ-H2AX* (magenta) and graphical representation of *ɣ-H2AX* positive cells undergoing DNA damage upon either PSC niche- specific (using *collierGal4*, Fig. S5D) or medullary zone hematopoietic progenitor- specific (using *domeGal4*, Fig. S5H) or cortical zone differentiated hemocytes- specific (using *hmlΔGal4*, Fig. S5L) or whole lymph- specific (using *e33cGal4*, Fig. S5P) or systemic fat body- specific (using *pplGal4*, Fig. S5T) expression of *UAS-foxoGFP* (Fig. S5B - R) or *UAS-pirkRNAi* (Fig. S5C - S) as compared to respective *wildtype* control (Fig. S5A - Q). GFP expression (Green) is either driven by *collierGal4* (Fig. S5A - C) or *domeGal4* (Fig. S5E - G) or *hmlΔGal4* (Fig. S5I - K). Nuclei are stained with DAPI (Blue). For PSC niche - mediated *ɣ-H2AX* positive cells undergoing DNA damage, *collierGal4 x UAS-foxoGFP* (N = 13, n = 25) or *collierGal4 x UAS-pirkRNAi* (N = 9, n = 18) were analyzed as compared to *collierGal4 x wt* (N = 17, n = 34). For hematopoietic progenitor mediated *ɣ-H2AX* positive cells, *domeGal4 x UAS-foxoGFP* (N = 11, n = 22) or *domeGal4 x UAS-pirkRNAi* (N = 12, n = 24) were analyzed as compared to *domeGal4 x wt* (N = 16, n = 32). For differentiated hemocyte - mediated *ɣ-H2AX* positive cells, *hmlΔGal4 x UAS-foxoGFP* (N = 14, n = 27) or *hmlΔGal4 x UAS-pirkRNAi* (N = 24, n = 47) were analyzed as compared to *hmlΔGal4 x wt* (N = 13, n = 26). For whole lymph gland – mediated *ɣ-H2AX* positive cells, *e33cGal4 x UAS-foxoGFP* (N = 12, n = 23) or *e33cGal4 x UAS-pirkRNAi* (N = 23, n = 45) were analyzed as compared to *e33cGal4 x wt* (N = 13, n = 25). For fat body – mediated *ɣ-H2AX* positive cells, *pplGal4 x UAS-foxoGFP* (N = 14, n = 28) or *pplGal4 x UAS-pirkRNAi* (N = 14, n = 28) were analyzed as compared to *pplGal4 x wt* (N = 17, n = 33). N denotes the number of larvae & n denotes the number of individual primary lymph gland lobes analyzed per genotype. Individual data points in the graphs represent individual primary lobes of the Lymph gland. Values are mean ± SD, and asterisks denote statistically significant differences (ns denotes not significant, **p < 0.01, *** p < 0.001, ****p < 0.0001). Student's t-test with Welch's correction was performed for the statistical analysis. Scale Bar: 50 µm (A-S).

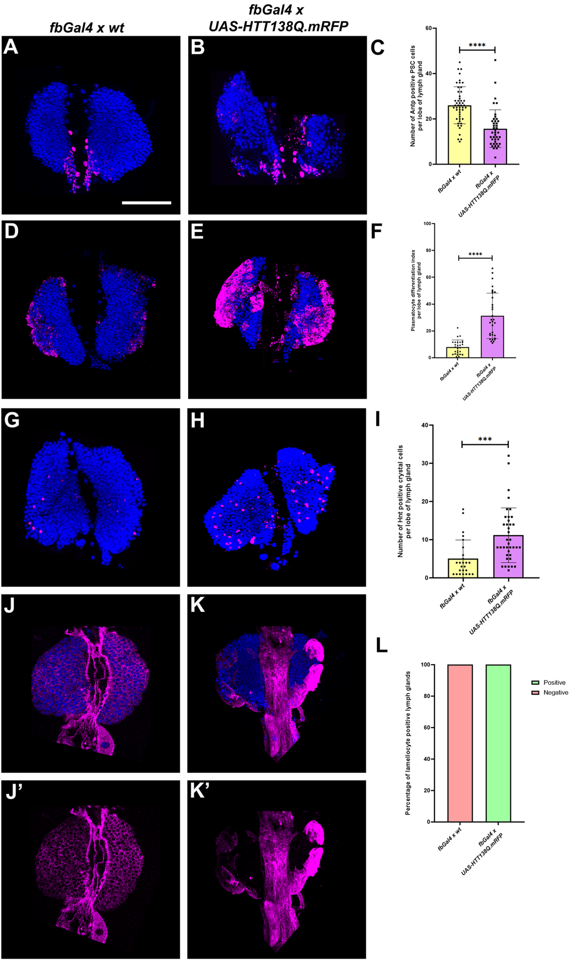
fbGal4 x wt
fbGal4 x
UAS-HTT138Q.mRFP
A
B
C
D
E
F
G
H
I
J
K
L
J'
K'
DAPI Antp
DAPI P1
DAPI Hnt
DAPI Mys
Mys
Number of Antp positive PSC cells per lobe of lymph gland
****
Plasmatocyte differentiation index per lobe of lymph gland
****
Number of Hnt positive crystal cells per lobe of lymph gland
***
Percentage of lamellocyte positive lymph glands
Positive
Negative
fbGal4 x wt
fbGal4 x UAS-HTT138Q.mRFP
0
20
40
60
80
100
10
30

**Supplementary Figure 6: Genetic induction of proteotoxic stress in fat body systemically affects lymph gland homeostasis in *Drosophila***

Posterior Signalling Centre (PSC) niche cell population marked by Antennapedia (Antp in magenta, Fig. S6B) or Plasmatocyte differentiation marked by P1 (magenta, Fig. S6E) or Crystal cell numbers marked by Hindsight (Hnt in magenta, Fig. S6H) or Lamellocyte differentiation marked by β-integrin specific Myospheroid (Mys in magenta, Fig. S6K-K') upon genetic induction of proteotoxic stress by expression of *HTT*138Q.m*RFP* in the fat body (using *fbgal4*) as compared to respective *wildtype* control (Fig. S6A, D, G, J-J'). Nuclei are stained with DAPI (Blue). Graphical representation of PSC niche numbers (Fig. S6C) or plasmatocyte differentiation index (Fig. S6F) or Crystal cell numbers (Fig. S6I) or percentage of lamellocyte positive lymph glands (Fig. S6L) upon genetic induction of proteotoxic stress in the fat body as compared to their respective *wildtype* control. For genetic induction of proteotoxic stress in fat body, the PSC niche numbers (*fbGal4 x wt* – N = 25, n = 49; *fbGal4 x UAS-HTT138Q.mRFP* - N = 21, n = 41), Plasmatocyte differentiation (*fbGal4 x wt* – N = 14, n = 28; *fbGal4 x UAS-HTT138Q.mRFP* - N = 16, n = 31), Crystal cell numbers (*fbGal4 x wt* – N = 13, n = 25; *fbGal4 x UAS-HTT138Q.mRFP* - N = 20, n = 40) and lamellocyte differentiation (*fbGal4 x wt* – N = 19; *fbGal4 x UAS-HTT138Q.mRFP* - N = 17) were analyzed per genotype. N denotes the number of larvae & n denotes the number of individual primary lymph gland lobes analysed per genotype. Individual data points in the graphs represent individual primary lobes of the Lymph Gland. Values are mean ± SD, and asterisks denote statistically significant differences (ns denotes not significant, $^{**}p < 0.01$, $^{***}p < 0.001$, $^{****}p < 0.0001$). Student's t-test with Welch's correction was performed for the statistical analysis. Scale Bar: 50 µm (A-K').

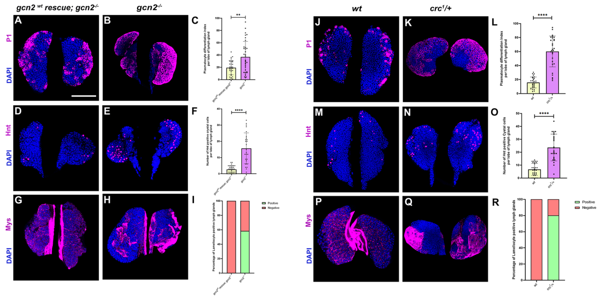

gcn2 wt rescue; gcn2-/-
gcn2-/-
wt
crc1/+
A
B
C
D
E
F
G
H
I
J
K
L
M
N
O
P
Q
R
P1
DAPI
Hnt
Mys
Plasmatocyte differentiation index per lobe of lymph gland
Number of Hnt positive crystal cells per lobe of lymph gland
Percentage of Lamellocyte positive lymph glands
Number of Hnt positive Cystal cells per lobe of lymph gland
Positive
Negative
**
****
wt

**Supplementary Figure 7: ISR pathway mutants exhibit defective lymph gland homeostasis**

Plasmatocyte differentiation marked by P1 (magenta, Fig. S7B, K) or crystal cell numbers marked by Hindsight (Hnt in magenta, Fig. S7E, N) or Lamellocyte differentiation marked by β-integrin specific Myospheroid (Mys in magenta, Fig. S7H, Q) in whole animal $gcn2^{-/-}$ homozygous null mutant or $crc^{1/+}$ heterozygous hypomorphic mutant LGs as compared to $gcn2^{wt}$ rescue; $gcn2^{-/-}$ (Fig. S7A, D, G) or *wildtype* (Fig. S7J, M, P) controls respectively. Graphical representation of Plasmatocyte differentiation index (Fig. S7C, L) or Crystal cell numbers (Fig. S7F, O) or percentage of lamellocyte positive lymph glands (Fig. S7I, R) in whole animal $gcn2^{-/-}$ null mutant or $crc^{1/+}$ heterozygous mutant as compared to the respective $gcn2^{wt}$ rescue; $gcn2^{-/-}$ (Fig. S7C, F, I) or *wildtype* (Fig. S7L, O, R) controls. $gcn2^{-/-}$ *and* $crc^{1/+}$ mutants for Plasmatocyte differentiation ($gcn2^{wt}$ rescue; $gcn2^{-/-}$ - N = 17, n = 34; $gcn2^{-/-}$ - *N* = 17, *n = 34 and wt* – N = 15, n = 29*;* $crc^{1/+}$ - N = 14, n = 28) or crystal cell numbers ($gcn2^{wt}$ rescue; $gcn2^{-/-}$ - N = 18, n = 35; $gcn2^{-/-}$ - *N* = 14, *n* = 27 *and wt – N* = 15, n = 29*;* $crc^{1/+}$ - N = 12, n = 24) or percentage of lamellocyte positive lymph glands ($gcn2^{wt}$ rescue; $gcn2^{-/-}$ - N = 15; $gcn2^{-/-}$ - *N* = 15 *and wt – N* = 15*;* $crc^{1/+}$ - N = 15) were analyzed per genotype. N denotes the number of larvae & n denotes the number of individual primary lymph gland lobes analyzed per genotype. Individual data points in the graphs represent individual primary lobes of the Lymph Gland. Values are mean ± SD, and asterisks denote statistically significant differences (ns denotes not significant, **$p < 0.01$, *** $p < 0.001$, ****$p < 0.0001$). Student's t-test with Welch's correction was performed for the statistical analysis. Scale Bar: 50 µm (A-Q).

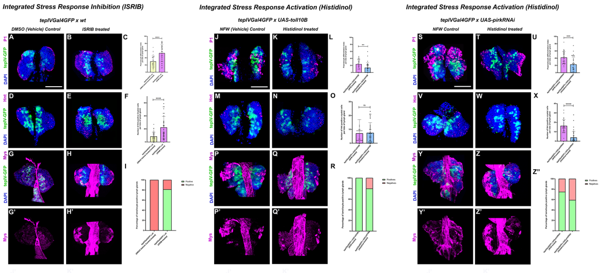

Integrated Stress Response Inhibition (ISRIB)
tepIVGal4GFP x wt
DMSO (Vehicle) Control
ISRIB treated
Integrated Stress Response Activation (Histidinol)
tepIVGal4GFP x UAS-toll10B
NFW (Vehicle) Control
Histidinol treated
Integrated Stress Response Activation (Histidinol)
tepIVGal4GFP x UAS-pirkRNAi
NFW Control
Histidinol treated
P1
tepIV-GFP
DAPI
Hnt
Mys
A
B
C
D
E
F
G
H
I
G'
H'
J
K
L
M
N
O
P
Q
R
P'
Q'
S
T
U
V
W
X
Y
Z
Z''
Y'
Z'
Positive
Negative
Positives
Negatives
Percentage of lamellocyte positive lymph glands

**Supplementary Figure 8: Chemical modulation of ISR pathway regulates lymph gland homeostasis in *Drosophila***

Plasmatocyte differentiation marked by P1 (magenta) or Crystal cell differentiation marked by Hindsight (Hnt in magenta) or lamellocyte differentiation marked by Myospheroid (Mys in magenta) in *tepIVGal4 x wt* larvae (Fig. S8B, E, H – H') upon ISR inhibition by ISRIB treatment as compared with DMSO (vehicle) control (Fig. S8A, D, G - G') or *tepIVGal4 x UAS-toll10B* (Fig. S8K, N, Q – Q') *or tepIVGal4 x UAS-pirkRNAi (*Fig. S8T, W, Z – Z'*)* larvae upon ISR activation by treatment with Histidinol as compared with respective NFW (vehicle) control (Fig. S8J, M, P – P', S, V, Y – Y'). Graphical representation of Plasmatocyte differentiation index (Fig. S8C, L, U) or number of Crystal cells (Fig. S8F, O, X) or percentage of Lamellocyte positive lymph glands (Fig. S8I, R, Z") in *tepIVGal4 x wt* larvae upon ISR inhibition by ISRIB treatment (Plasmatocytes: N = 16, n = 32; Crystal cells: N = 17, n = 33; Lamellocytes: N = 16) as compared with the respective vehicle control (Plasmatocytes: N = 16, n = 32; Crystal cells: N = 16, n = 32; Lamellocytes: N = 18) or *tepIVGal4 x UAS-toll10B* larvae upon ISR activation by Histidinol treatment (Plasmatocytes: N = 15, n = 30; Crystal cells: N = 16, n = 32; Lamellocytes: N = 17) as compared to the respective vehicle control (Plasmatocytes: N = 15, n = 30; Crystal cells: N = 15, n = 30; Lamellocytes: N = 16) *or tepIVGal4 x UAS-pirkRNAi* larvae upon ISR activation by Histidinol treatment (Plasmatocytes: N = 16, n = 32; Crystal cells: N = 18, n = 36; Lamellocytes: N = 32) were analyzed as compared to the respective vehicle control (Plasmatocytes: N = 15, n = 30; Crystal cells: N = 13, n = 26; Lamellocytes: N = 12). GFP (green) is driven by *tepIVGal4* which marks core progenitors in the LG (S8A-Z'). Nuclei are stained with DAPI (Blue). N denotes the number of larvae & n denotes the number of individual primary lymph gland lobes analysed per genotype. Individual data points in the graphs represent individual primary lobes of the Lymph Gland. Values are mean ± SD, and asterisks denote statistically significant differences (ns denotes not significant, $**p < 0.01$, $*** p < 0.001$, $****p < 0.0001$). Student's t-test with Welch's correction was performed for the statistical analysis. Scale Bar: 50 µm (A-Z').
